# Effects of climatic warming on spring phenology in subtropical trees: process-based modelling with experiments designed for model development

**DOI:** 10.1101/2020.12.29.422625

**Authors:** Rui Zhang, Jianhong Lin, Fucheng Wang, Heikki Hänninen, Jiasheng Wu

**Author notes:** R. Zhang and J. Lin are to be considered joint first authors. **Correspondence** Jiasheng Wu, Heikki Hänninen. **Funding information**, The study was financed by The Chinese National Natural Science Foundation [31800579], The National Forestry and Grassland Technological Innovation Program for Young TopNotch Talents [2020132604], Zhejiang Provincial Natural Science Foundation of China [LQ18C160001], the Key Research Program of Zhejiang Province [2018C02004], the Major Project for Agricultural Breeding of Zhejiang Province [2016C02052-12], and Overseas Expertise Introduction Project for Discipline Innovation (111 Project D18008).

## Abstract

To project the effects of climatic warming on the timing of spring leafout and flowering in trees, process-based tree phenology models are often used nowadays. Unfortunately, the biological realism of the models is often compromised because the model development has often been based on various assumptions and indirect methods. We developed process-based tree phenology models for four subtropical tree species, and for the first time for these trees, we based the model development on explicit experimental work particularly designed to address the processes being modelled. For all the four species, a model of seedling leafout was developed, and for *Torreya grandis*, a model for female flowering in adult trees was additionally developed. The models generally showed reasonable accuracy when tested against two sources of independent data: observational phenological records and leafout data from a whole-tree chamber warming experiment. In scenario simulations, the models projected an advanced spring phenology under climatic warming for 2020 – 2100. For the leafout of seedlings, the advancing rates varied between 4.7 and 5.9 days per one °C warming, with no major differences found between the climatic scenarios RCP4.5 and RCP8.5. For *Torreya* flowering, less advancing was projected, and the projected advancing per one °C warming was less for RCP8.5 (0.9 days / °C) than for RCP4.5 (2.3 days / °C). The low advancing rates of *Torreya* flowering were caused by reduced chilling under the warming climate and by the particular temperature responses found for *Torreya* flowering. For instance, our results show that in *Torreya* flower buds, no rest break (endodormancy release) is seen at +15 °C, whereas in the seedlings of all four species, +15 °C has a clear rest-breaking effect. These findings highlight the need to base the model development on explicit experiments particularly designed to address the process being modelled.

## 1 Introduction

In the terrestrial biomes of the world, shifts in the spring phenology of plants are among the most important effects of the ongoing climate change (Polgar & Primack, 2011). These changes affect essential ecosystem processes, such as the cycling of water, carbon and nutrients (Kramer & Hänninen, 2009; Richardson et al., 2009), ecosystem productivity (Keenan et al., 2014), plant-animal relationships (Senior, Evans, Leather, Oliver, & Evans, 2020), population dynamics and competition of plant and animal species (Delpierre, Guillemot, Dufrêne, Cecchini, & Nicolas, 2017; Zettlemoyer, Schultheis, & Lau, 2019), and ultimately, shifts in the geographical ranges of species (Chuine & Beaubien, 2001; Chuine, 2010). Analyses of long-term phenological records have generally shown that warming has caused advanced phenological development in spring over the past few decades (Menzel et al., 2006; Zheng et al., 2016). However, the sensitivity of the phenological development to rising temperatures has been found to decrease at the same time, so that the advancing trends of spring phenology have been levelling off in several cases (Fu et al., 2015a). This has been attributed to the effects of reduced chilling, also caused by climatic warming (Murray, Cannell, & Smith, 1989; Fu et al., 2015b; Ford, Harrington, Bansal, Gould, & St Clair, 2016; Luedeling, 2012; Martínez-Lüscher, Hadley, Ordidge, Xu, & Luedeling, 2017, Wang et al., 2020;), or to the restricting effect of photoperiod, which is not changed by climatic warming (Vitasse & Basler, 2013; Fu et al., 2015a, 2019a,b).

After growth cessation in autumn, the buds of boreal and temperate trees attain the state of rest (endodormancy; Lang, Early, Martin, & Darnell, 1987), where bud burst and growth onset are prevented by physiological factors inside the buds (Perry, 1971; Fuchigami, Weiser, Kobayashi, Timmis, & Gusta, 1982; Hänninen & Tanino, 2011). The growth-arresting conditions are removed by prolonged (days to months) exposure to low chilling temperatures; traditionally, temperatures near +5 °C have been regarded as necessary for chilling (Sarvas, 1974). This is the chilling requirement of rest completion (Hänninen, 1995, 2016), which prevents a premature growth onset during mild periods in autumn and winter, which would lead to frost damage during subsequent periods of frost (Cannell, 1985; Hänninen, 1991; Zohner, Mo, Sebald, & Renner, 2020), a phenomenon more recently called ‘false springs’ (Marino, Kaiser, Gu, & Ricciuto, 2011; Chamberlain, Cook, de Cortázar-Atauri, & Wolkovich, 2019). After the chilling requirement is met, the phase of quiescence (ecodormancy, Lang et al, 1987) is attained (Perry, 1971; Fuchigami et al., 1982; Hänninen & Tanino, 2011). During quiescence, bud burst and growth onset are prevented by low air temperatures only, so that ontogenetic development, i.e., microscopic anatomic changes within the buds towards bud burst, occurs in relatively high growth-promoting forcing temperatures (Sarvas 1972, 1974). Visible bud burst occurs as a result of prolonged (days to months) exposure to forcing temperatures (the high-temperature requirement of growth onset, Hänninen, 1995, 2016).

The effects of high temperatures on the spring phenology of plants have been addressed in quantitative terms with various temperature sum calculations since the days of Réaumur (1735), (for reviews, see Arnold, 1959; Wang, 1960; Chuine, de Cortázar-Atauri, Kramer, & Hänninen, 2013). Similar chilling-unit calculations for rest break are of more recent origin, but they, too, have a long history by now (Weinberger, 1950). Since the early 1970s, holistic models addressing both phenomena simultaneously have been developed for boreal and temperate trees (Sarvas, 1972, 1974; Richardson et al., 1974; Landsberg, 1974; Chuine et al., 2016; Harrington, Gould, & St.Clair, 2010). These models, currently known as process-based tree phenology models, predict the timing of spring phenological events, such as vegetative bud burst or flowering, on the basis of air temperature data (Hänninen & Kramer, 2007; Chuine et al., 2013). Sometimes these models also address the effects of photoperiod (Nizinski & Saugier, 1988; Hänninen, 1995). Since the mid-1980s, process-based tree phenology models have been increasingly used for assessing the effects of the projected climate change (Cannell, 1985; Hänninen, 1991; Kramer, 1994; Chuine et al., 2016).

In boreal and temperate trees, the chilling requirement of rest break has been studied for a hundred years: the phenomenon was first reported by Coville (1920). Given this background, it is surprising that the rest period and the chilling requirement in subtropical trees have not been experimentally examined until recently, when Du, Pan, & Ma (2019), Song et al. (2020), and Zhang et al. (2020) showed that subtropical trees also exhibit the rest period and the chilling requirement of rest completion. Further, Zhang et al. (2021) have recently shown that the air temperature responses of the dormancy processes in subtropical trees are different from those in the more northern ones. This points to the importance of developing specific process-based tree phenology models for subtropical trees. To our knowledge, no process-based model has been developed for any subtropical tree species by using data from experiments explicitly designed for developing and parameterising the model. Here, using the experimental results of Zhang et al. (2021) and some additional experimental data gathered for the present study, we developed process-based spring phenology models for four subtropical tree species. The models were tested with three sources of independent data and then used in scenario simulations to assess the effects of climatic warming on spring phenology in the four sub-tropical tree species over the years 2020 – 2100. We hypothesised that due to differences in the dormancy processes addressed by the models, the different species (Zhang et al., 2021), different life stages (Vitasse, 2013), and different bud types of a given species (Ma, Huang, Hänninen, Li, & Berninger, 2020) would respond to the warming differentially. The models we develop here will facilitate the assessing of the effects of climatic change in subtropical conditions.

## 2 MATERIALS AND METHODS

### 2.1 Model structure

The process-based tree phenology models developed in the present study simulate the effects of air temperature on the processes of rest break and microscopic ontogenetic development in the buds towards bud burst (Figure 1). Each model developed was constructed as a combination of three sub-models: one for rest break, one for ontogenetic development, and one for mediating the effects of rest break on the ontogenetic development. This modular model structure facilitates the use of experimental designs that explicitly address each of the three modelled physiological phenomena at the whole-tree level (Hänninen et al., 2019).

**FIGURE 1.**
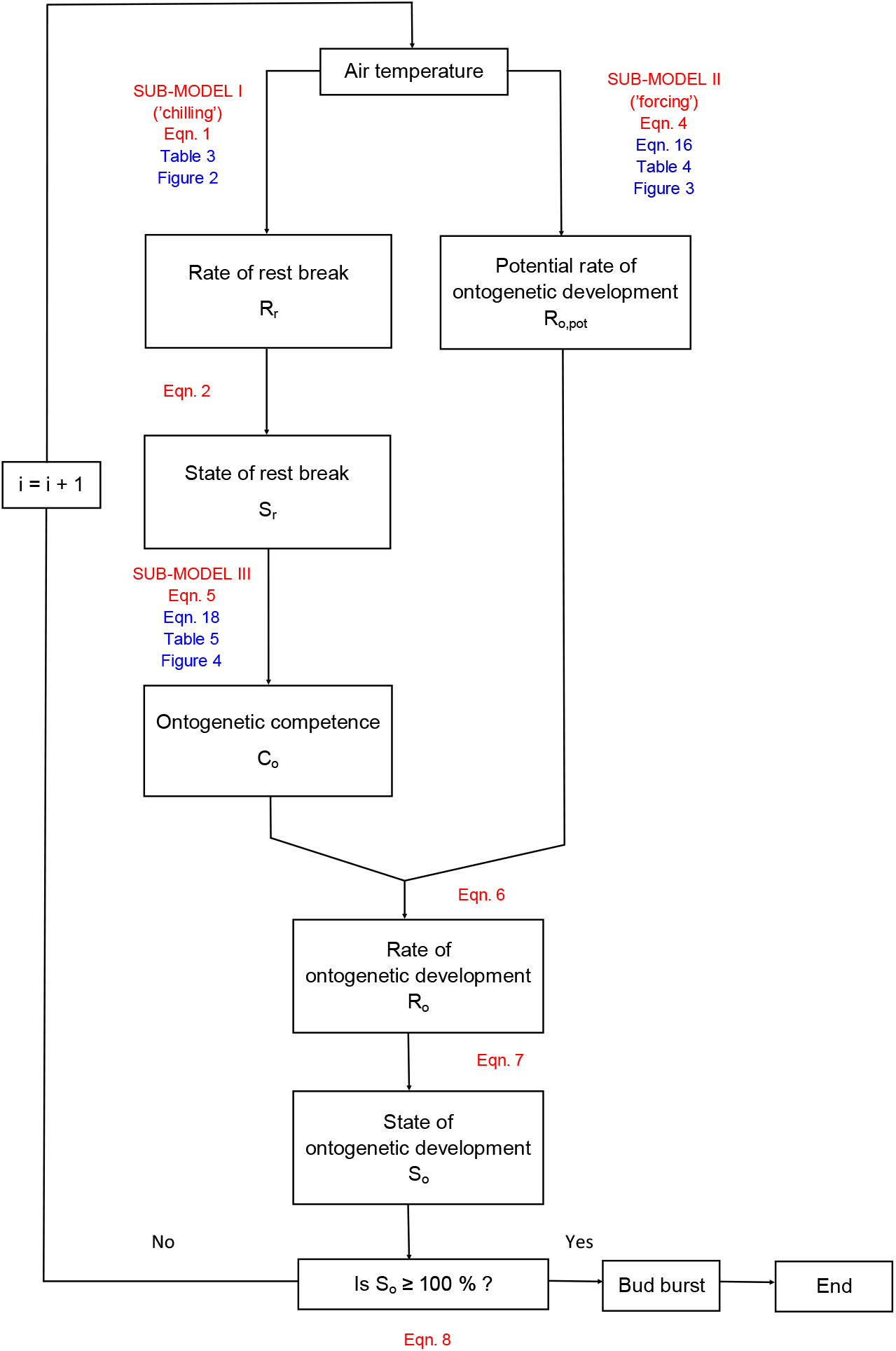
The modular model structure applied in the present study for process-based modelling of bud burst phenology in four subtropical tree species (Hänninen, 1990, 2016; Hänninen & Kramer, 2007). The overall model consists of three sub-models, each one addressing the corresponding particular ecophysiological phenomenon that affects the timing of bud burst. The red font indicates the general structure of the overall model and the blue font the specific models formulated in the present study on the basis of experimental data. In the present study, the model was applied with a time step of one hour, using the hourly air temperature as input in the simulations. Modified from Hänninen (2016)

Sub-model I addresses the effects of low chilling temperatures on the process of rest break (Figure 1). The momentary rate of rest break, R_r_(t), is first calculated as a function of air temperature, T(t):

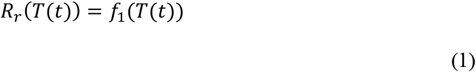

where f_1_ = the function to be determined on the basis of experimental data (see Section 2.3.2). By definition, the state of rest break at time instant t, S_r_(t), is obtained by integrating the momentary values of the rate of rest break, R_r_(T(t)), with respect to time:

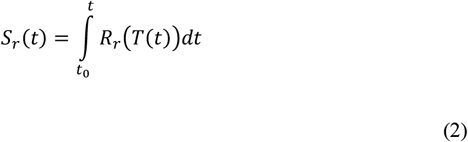

where to = the starting moment of the rest period. In practice the integration is carried out by summing the hourly values of R_r_(T(i)) calculated on the basis of hourly temperatures, T(i). Rest completion is predicted to occur when the value of S_r_(t) attains 100 %:

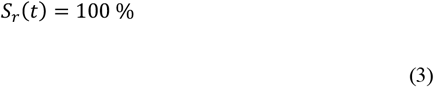

After rest completion, the value of S_r_(t) is set at 100 % for the remaining simulation period, which ends at the predicted time of bud burst.

Sub-model II addresses the effects of high forcing temperatures on the rate of ontogenetic development after rest completion. That particular rate is called the potential rate of ontogenetic development, R_o,pot_, because after rest completion the rate of ontogenetic development is no longer restricted by the rest condition. The momentary R_o,pot_ is calculated as a function of air temperature, T(t):

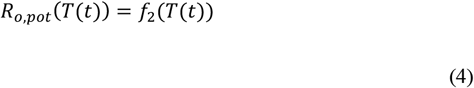

where f_2_ = the function to be determined on the basis of experimental data (see Section 2.3.3).

Sub-model III addresses the restricting effects of the bud rest status on the rate of ontogenetic development. These effects are modelled with the aid of the model variable ontogenetic competence, C_o_, which is a multiplier with values ranging from 0 to 1. With C_o_ = 0, there is no ontogenetic development regardless of the prevailing temperature. At the time of rest completion, the value of C_o_ = 1 is attained, and the rate of ontogenetic development is then equal to the potential rate. With values between 0 and 1, ontogenetic development takes place at a reduced rate, as indicated by the decimal of C_o_. Ontogenetic competence, C_o_, mediates the effects of the rest status onto the ontogenetic development, so that its momentary value is calculated as a function of the state of rest break, S_r_:

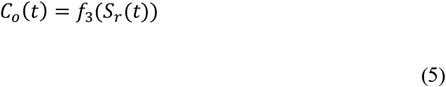

where f_3_ = the function to be determined on the basis of experimental data (see Section 2.3.4).

Following the definition of the variable ontogenetic competence, C_o_, the momentary rate of ontogenetic development, R_o_(t) is calculated as

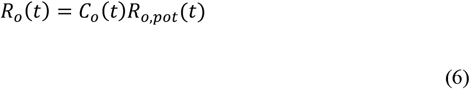

By definition, the state of ontogenetic development at time instant t, S_o_(t), is obtained by integrating the momentary value of the rate of ontogenetic development, R_o_(T(t)), with respect to time:

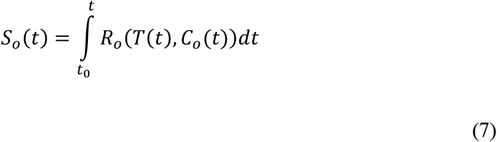

where t_o_ = the starting moment of the rest period. In practice the integration is carried out by summing the hourly values of R_o_(T(i)) calculated on the basis of the hourly temperature, T(i). Bud burst is predicted to occur, and the simulation ends, when the value of S_r_(t) attains 100 %:

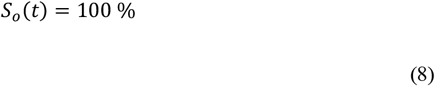

### 2.2 Tree species and experimental data

#### 2.2.1 Material categories

Process-based tree phenology models were constructed for four sub-tropical tree species: *Castanopsis sclerophylla, Phoebe chekiangensis, Pseudolarix amabilis*, and *Torreya grandis*. For all species, first-year container seedlings were used in the experiments. With *Torreya*, experiments were also carried out with twigs detached from adult trees. Accordingly, a total of five process-based models, each representing the corresponding material category (species, seedling / adult tree), were constructed. To develop the models, both previously published (Autumn experiments, Zhang et al., 2021) and original (Spring experiments) data were used.

#### 2.2.2 Autumn experiments for Sub-models I and III

In the autumn experiments carried out in 2017 – 2019, the experimental seedlings and twigs were first subjected to chilling for periods of varying duration, then transferred to growth-promoting forcing conditions with a constant air temperature of +20 °C (Zhang et al., 2021). The chilling treatments were carried out in growth chambers at three constant temperatures: +5 °C, +10 °C, and +15 °C. In the forcing conditions, a regrowth test was carried out by observing the occurrence and timing of bud burst (leafout, flowering; see Section 2.2.3). For each observed seedling or twig showing bud burst, the days to bud burst in the forcing conditions, DBB, was calculated. For each treatment representing a given duration of chilling at a given chilling temperature, the value of bud burst percentage, BB%, was calculated.

#### 2.2.3 Spring experiments for Sub-model II

The effects of the air temperature on the potential rate of ontogenetic development were addressed in the three spring experiments, carried out in the years 2018 – 2020. Until the start of the experiment, the experimental seedlings and trees overwintered in natural conditions on the Zhejiang A&F University campus (30°14′N, 119°42′E) in Hangzhou, southeastern China. In order to make sure that the actual rate of ontogenetic development observed in the experiment represented the potential rate, the spring experiments of all the three years 2018 – 2020 were started on 13 February, after all the material categories had met the chilling requirement in natural conditions (Zhang et al., 2021). In each year, we transferred ten replicated seedlings of each species from natural outdoor conditions to forcing growth chambers (E-Lotus Technology Co., Beijing, China) with several constant air temperatures ranging from 10 to 28 *°C* (Table 1). At the same time, twigs of adult *Torreya* trees were sampled on the campus and transferred to the forcing chambers. The twigs and seedlings were prepared for the experiments in the same way as in the autumn experiments (Zhang et al., 2021). In all the forcing chambers, day length was 12 h, PPFD 400 μmol m^−2^ s^−1^, relative humidity 70-80 %, and concentration of CO_2_ 300-400 ppm. Temperature of the chambers was monitored with iButton (Model DS1912L, Embedded Data Systems Co., Ltd, KY, USA). Bud burst was observed visually, as had been done in the autumn experiments (Zhang et al., 2021). In the seedlings of all four species, the determination of bud burst was based on the stage of leafout, and in the *Torreya* twigs representing adult trees, on female flowering. Despite these differences, the term ‘bud burst’ is used in this report as a generic one in connection with the standard variables BB% and DBB (Zhang et al., 2021). For each experimental seedling and twig, the days to bud burst in the forcing conditions, DBB, was calculated.

**Table 1.** Experimental temperatures applied in the Spring experiments of 2018 – 2020 for formulating Sub-model II for each of the five material categories addressed in the study (seedlings of three species and both seedlings and twigs representing adult trees of *Torreya*).

| Year | Experimental temperature |
| --- | --- |
| 2018 | +10 °C |
|  | +17 °C |
|  | +24 °C |
| 2019 | +12 °C |
|  | +20 °C |
|  | +28 °C |
| 2020 | +10 °C |
|  | +15 °C |
|  | +20 °C |
|  | +25 °C |

### 2.3 Formulating the models on the basis of experimental data

#### 2.3.1 Outline

The experimental information was introduced to Sub-models I, II, and III via the functions f_1_ (Eqn. 1), f_2_ (Eqn. 4), and f_3_ (Eqn. 5), respectively. Once these functions had been defined and parameterized on the basis of the experimental data, the overall model defined by Eqns. (1) – (8) was operationalized to be used in computer simulations.

#### 2.3.2 Sub-model I

The rate of rest break at any constant temperature T’, R_r_(T’) was determined experimentally as follows (Sarvas, 1974):

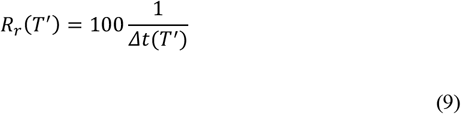

where Δt(T’) = the duration of chilling required for rest completion (meeting the chilling requirement) at the constant temperature T’. Because of the multiplier 100, the unit of R is % h^−1^, so that the value of R_r_(T’) indicates how many per cent of the cumulative physiological processes required for rest completion take place within one hour. The value of Δt was determined on the basis of the results of the autumn experiments. First, the values of bud burst percentage, BB%, and days to bud burst, DBB, were plotted against the duration of chilling. By means of the scatter plots thus obtained, the value of Δt needed in Equation (9) was determined by two rules: First, it was required that BB% ≥ 80 % for the chilling duration corresponding to Δt. Second, it was required that the value of DBB decreased to a threshold value, DBB_RC_, determined by the levelling off of the DBB curve in the +5 °C chilling experiment. To that end, an exponential function f4 was first fitted to the scatter plot of the +5 °C chilling experiment, representing the DBB value of each seedling/twig as a function of the duration of chilling (Figure S1a):

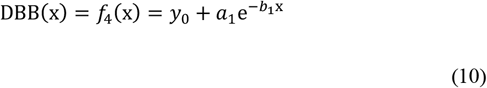

where DBB is the modelled DBB, x = the duration of chilling (days); and y_o_, a_1_, and b_1_ are parameters to be estimated. Next, the first derivative with respect to the duration of chilling of the fitted function was calculated as follows:

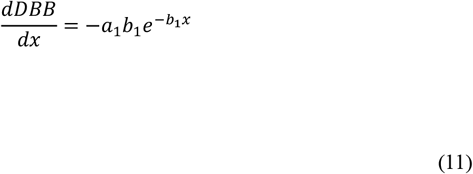

In the next step, Δt was determined for the +5 °C chilling treatment as equal to the duration of chilling x where the value of the first derivative is equal to −0.3 (Figure S1a):

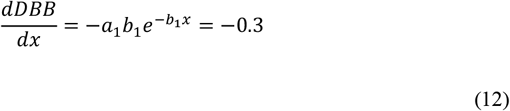

implying

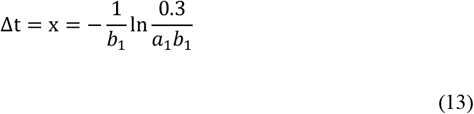

Next, the exponential function (Eqn. 10) was fitted to the scatter plots of both +10 °C (Figure S1b) and +15 °C chilling treatments. For them, the value of Δt was determined as the duration of chilling implying the same value of DBB = DBB_RF_ as was obtained with the +5 °C chilling treatment (Figure S1):

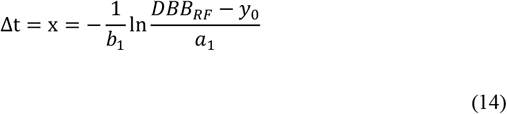

The Δt values obtained (Table 2; Figure S2) were plugged into Eqn. (9), thus getting three (for *Castanopsis, Phoebe*, and *Pseudolarix* seedlings) or four (for *Torreya* seedlings and twigs) data points for R_r_ in the temperature range of +5 to +15 °C (see Figure 2 in the Results Section). In that temperature range, the function f_1_ in Eqn. (1) was determined by fitting a direct line to the scatter plot representing the observed R_r_ as a function of temperature. It was assumed that at temperatures below +5 °C, R_r_ would drop from its maximum value (obtained at +5 °C) to zero at −3.4 °C (Sarvas, 1974). If extrapolation was also needed at the high temperature end, it was similarly assumed that R_r_ would drop from its value at +15 °C to zero at +20 °C. In a sensitivity analysis addressing the uncertainty caused by the lack of data for temperatures under +5 °C, a modified model was used. In that model it is assumed that at any temperature below +5 °C, R_r_ has a constant value, identical to the maximal value of R_r_ at +5 °C (see Figure 2 in the Results Section).

**FIGURE 2.**
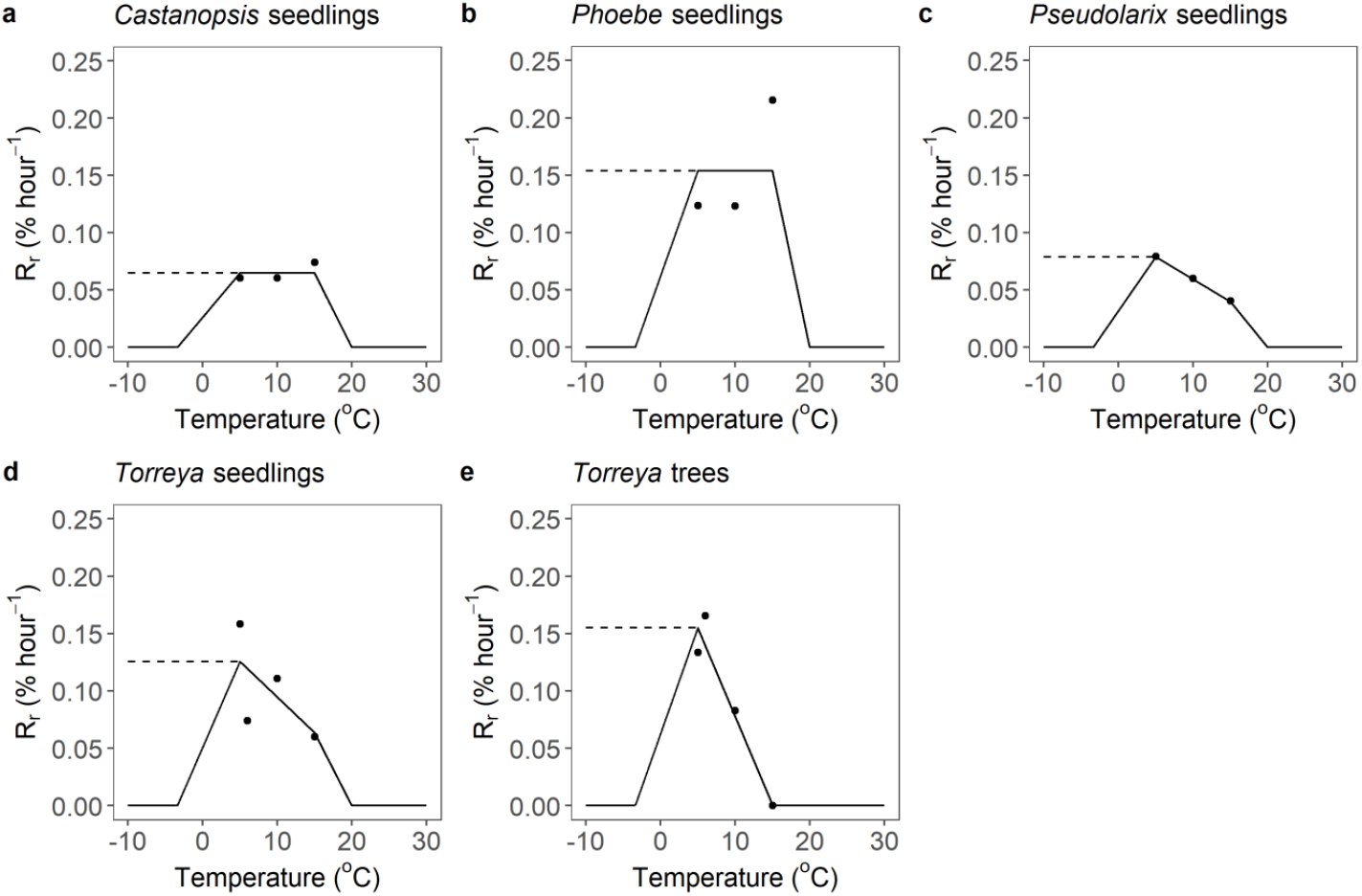
Sub-model I for each of the five material categories examined, representing the temperature response of the rate of rest break, R_r_, (in the overall model, function f_1_ in Eqn. 1). The data points were determined on the basis of the experimental results of Zhang et al. (2021). The equations for f_1_ are presented in Table 4. The value of the model variable R_r_ indicates how many per cent of the physiological changes required for rest completion take place in one hour. The value R_r_ = 0.1 % hour^−1^, for instance, implies that 1000 hours are required at the particular temperature for rest completion. The dotted lines indicate the modified model used in the sensitivity analysis that addressed the uncertainty caused by the lack of data for temperatures under +5 °C

**Table 2.** Duration of chilling, Δt, required at the experimental chilling temperatures for rest completion in the five material categories studied (seedlings of three species and both seedlings and twigs representing adult trees of *Torreya*). The values of Δt are indicated both in hours, as used in the modelling (Eqns. (1), (9); Table 4), and in days, as used in the plots to determine the values (Figures S1, Figure S2). No bud burst was observed in *Torreya* twigs after chilling at +15 °C.

| Material category | Chilling temperature | $\Delta t$, days | $\Delta t$, hours |
| --- | --- | --- | --- |
| *Castanopsis* seedlings | +5 °C | 68.70 | 1649 |
|  | +10 °C | 68.95 | 1655 |
|  | +15 °C | 56.25 | 1350 |
| *Phoebe* seedlings | +5 °C | 33.72 | 809 |
|  | +10 °C | 33.85 | 812 |
|  | +15 °C | 19.34 | 464 |
| *Pseudolarix* seedlings | +5 °C | 52.70 | 1265 |
|  | +10 °C | 69.45 | 1667 |
|  | +15 °C | 102.81 | 2467 |
| *Torreya* seedlings | +5 °C | 26.31 | 631 |
|  | +6 °C | 56.33 | 1352 |
|  | +10 °C | 37.56 | 901 |
|  | +15 °C | 69.51 | 1668 |
| *Torreya trees* | +5 °C | 31.21 | 749 |
|  | +6 °C | 25.20 | 605 |
|  | +10 °C | 56.00 | 1344 |
|  | +15 °C | - | - |

#### 2.3.3 Sub-model II

For Sub-model II, the potential rate of ontogenetic development at any constant temperature T’, R_o,pot_(T’), was determined experimentally as follows (Sarvas, 1972; Campbell, 1978):

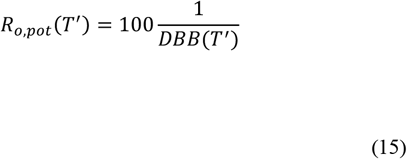

where DBB(T’) = days to bud burst measured at the constant temperature T’ after rest completion. Because of the multiplier 100, the unit of R_o,pot_ is % h^−1^, so that the value of R_r_(T’) indicates how many per cent of the ontogenetic changes required for bud burst take place within one hour.

The values of R_o,pot_(T’) were plotted against the experimental temperature T’. The air temperature response of R_o,pot_ (function f_2_ in Eqn. 4) was formulated by fitting a sigmoidal function to the scatter plot (see Figure 3 in the Results Section):

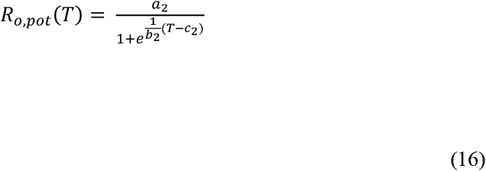

where T is air temperature and a_2_, and b_2_, and c_2_ are parameters to be estimated.

**FIGURE 3.**
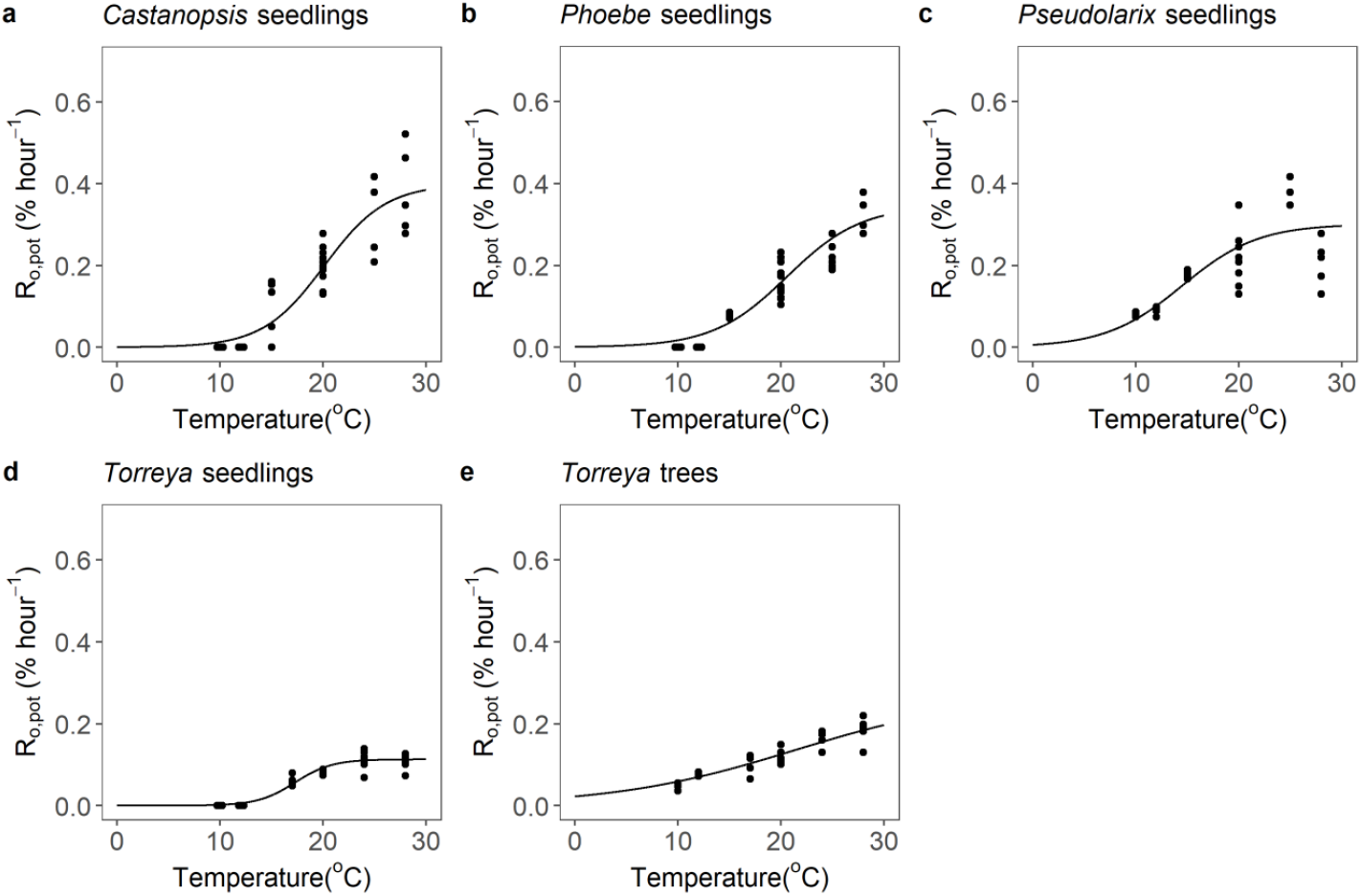
Sub-model II for each of the five material categories examined, representing the temperature response of the potential rate of ontogenetic development, R_o,pot_, (in the overall model, function f_2_ in Eqn. 4). Function f_2_ is formulated according to Eqn. (16) with the parameter values presented in Table 5. The potential rate of ontogenetic development indicates the rate at rest completion, i.e., when the chilling requirement has been met and the rate is no longer restricted by the rest condition of the bud. More specifically, R_o,pot_, indicates how many per cent of the ontogenetic changes within the bud required for leafout/flowering take place after rest completion in one hour. The value R_o,pot_ = 0.2 % hour^−1^, for instance, implies that at the particular temperature, 500 hours after rest completion are required for leafout/flowering

#### 2.3.4 Sub-model III

By the definition of ontogenetic competence, C_o_, (Section 2.1, Eqn. 6), the empirical value of C_o_ for a given seedling/twig is zero if it shows no bud burst during prolonged forcing, and by the same token, it is unity if its DBB is equal to the DBB representing twigs and seedlings which have just attained rest completion, i.e., equal to DBB(Δt) = DBB_RC_ (Figure S1). Between these two extremes, the value of C_o_ increases proportionally to the ratio of DBB(Δt) / DBB of the seedling/twig. In the calculation of the empirical values of C_o_, two averaging methods were applied. First, the original DBB observations of the individual twigs/seedlings were replaced by the fitted exponential functions, f4 (Eqn. 10; Figure S2). Accordingly, the value of C_o_ for any duration of chilling x at a given constant chilling temperature was calculated as the ratio f_4_(Δt) /f_4_(x). Second, in order to account for the seedlings/twigs showing little or no bud burst, the value of C_o_ was set at zero for chilling treatments in which fewer than a half of the buds showed bud burst:

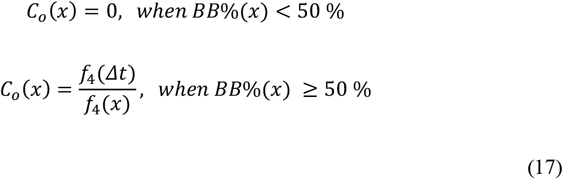

where C_o_ = ontogenetic competence, x = the duration of chilling, Δt = the duration of chilling required for rest completion, and BB% = bud burst percentage.

The value of ontogenetic competence, C_o_(t), was modelled with function f_3_ by using the state of rest break, S_r_(t), as the argument in f_3_ (Eqn. 5). In order to formulate function f_3_, the value of S_r_(x) was calculated for each duration of chilling x at a given constant chilling temperature as 100 x/Δt. This simple formula is implied in the assumption that at any constant air temperature, rest break progresses at a constant rate (Eqn. 9).

Next, the values of C_o_(x) were plotted against S_r_(x) for all constant chilling temperatures used in the experiment, and a piece-wise linear equation was fitted to the scatter plot of the data points (see Figure 4 in the Results Section):

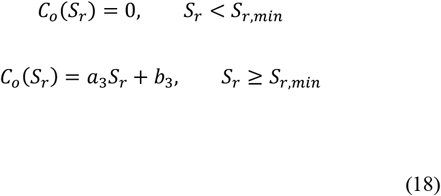

where C_o_ = ontogenetic competence, S_r_ = the state of rest break, S_r,min_ = the minimum value of S_r_ showing an above-zero value of C_o_ in the plot. The fitting was constrained by the assumption that the direct line crosses the point (100,1), since the value of Co(100) = 1 by definition. After rest completion at S_r_(t) = 100%, Eqn. (18) was not applied any longer; instead, the value of C_o_ = 1 was retained for the remaining simulation period, which ended in the predicted bud burst.

**FIGURE 4.**
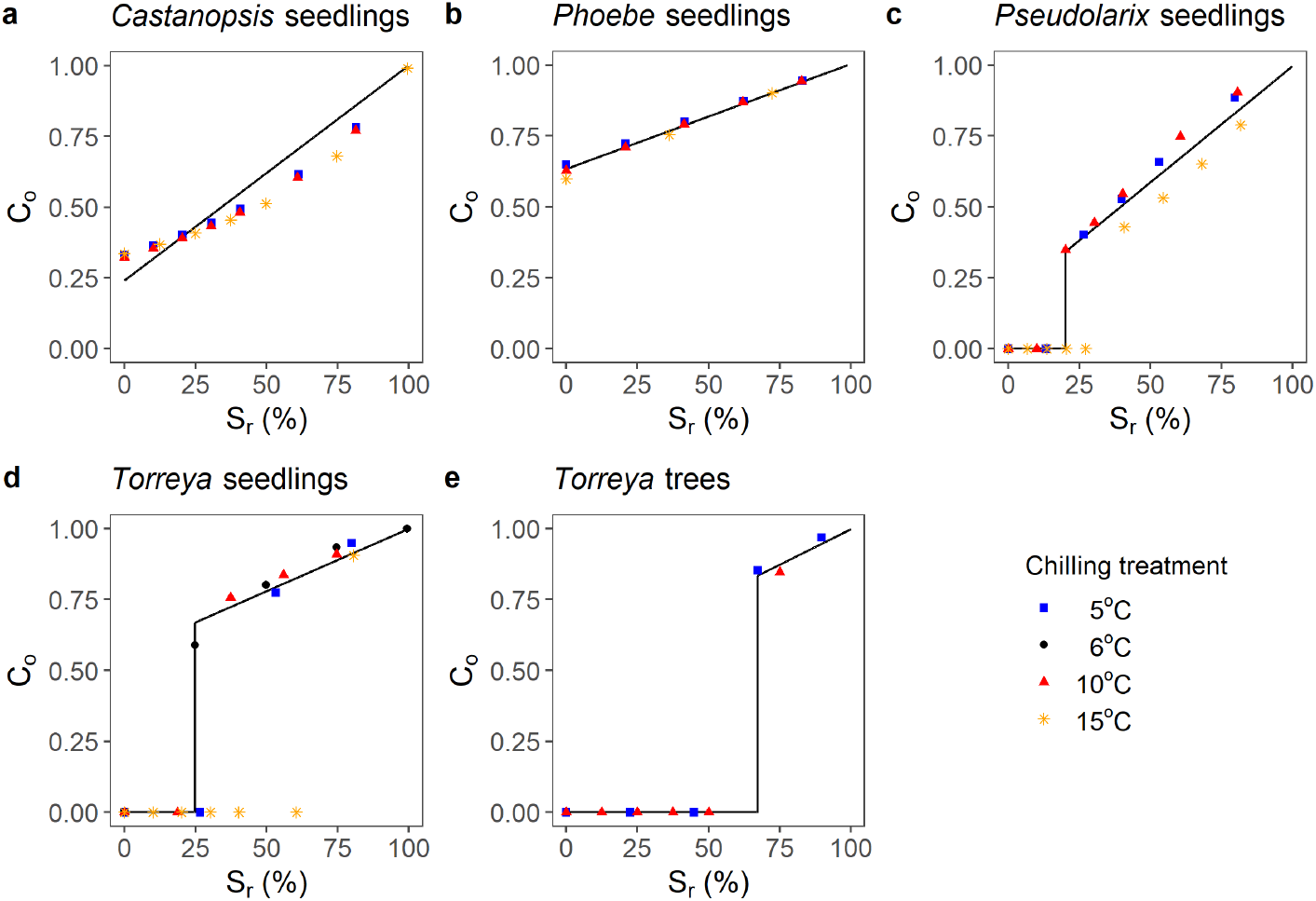
Sub-model IIII for each of the five material categories examined, representing the dependency of ontogenetic competence, C_o_, on the state of rest break, S_r_, (in the overall model, function f_3_ in Eqn. 5). The data points were determined on the basis of the experimental results of Zhang et al. (2021). Function f_3_ is formulated according to Eqn. (18) with the parameter values presented in Table 6. The value of S_r_ indicates the percentage of the physiological changes of rest break caused by chilling and required for rest completion that have taken place at a given moment. The value of C_o_ indicates the relative effect of the rest condition on the rate of ontogenetic development. With C_o_ = 0.5, for instance, twice as much time is required for leafout/flowering as is required after rest completion (Sr = 100 %) with C_o_ = 1

### 2.4 Model tests with independent bud burst data

#### 2.4.1 Testing procedure

For testing the models developed, three sets of independent bud burst observations were used. Starting on 23 November for each annual cycle, the model of each material category was run until it predicted bud burst to occur. Whenever hourly temperature data were not available, they were generated from the daily temperature indices as follows (Zohner et al., 2020):

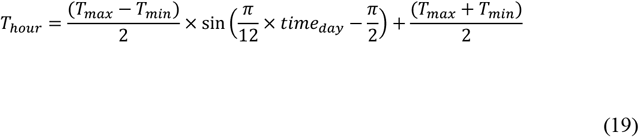

where T_min_ and T_max_ are the daily minimum and the daily maximum temperature, respectively, and time_day_ = the hour of the day addressed. The predicted bud burst dates were then compared with the observed ones.

#### 2.4.2 Predicted dormancy dynamics and a short-term independent test of the models

In order to examine the dynamics of rest break and ontogenetic development predicted by the models, we calculated the predicted state of rest break, S_r_, ontogenetic competence, C_o_, and the state of ontogenetic development, So, for the period of autumn 2018 to spring 2019 both in natural conditions on the Zhejiang A&F University campus and in natural conditions on a nearby *Torreya* tree plantation (Zhang et al., 2021). The calculations were based on hourly air temperature records collected by iButtons (Model DS1912L, Embedded Data Systems Co., Ltd, KY, USA) at the two respective sites. In order to provide the first short-term test with independent data, the timing of leafout predicted by the models was compared with leafout observations made in natural conditions on the campus with seedlings of all four species. Similarly, the timing of the flowering predicted for adult *Torreya* was compared with corresponding observations in the *Torreya* plantation.

#### 2.4.3 Phenology records

The models were tested against independent observational phenological records from the Chinese National Ecology Science Data Center (www.cnern.org.cn, Table 3). Within this network, phenological observations are made and recorded routinely by the professionals at each station in accordance with unified observation guidelines (China Meteorological Administration, 1993). Similarly to our experimental work, we determined vegetative bud burst on the basis of leafout and then, for testing the models, selected the date corresponding to it from the database. In addition, the phenological observations for *Torreya* flowering from Ren, Long, Zhou, Zhang, Zhou (2017) were used for independent model testing (Table 3). For the simulations, the daily T_min_ and T_max_ values were obtained from meteorological stations of the China National Meteorological Information Center (www.data.cma.cn) located near the phenological stations (Table 3). The values of T_min_ and T_max_ were converted into hourly temperatures by means of Eqn. (19). The tests for the three species other than *Torreya* were carried out with the model developed for seedlings because no models for adult trees had been developed for these species.

**Table 3.** The stations where phenological records were collected and the corresponding meteorological stations that provided air temperature data; both data sets were used in the present study for independent testing of process-based tree phenology models. For *Torreya*, the phenological data comprised female flowering (Station 4, Ren et al., 2019); for all other species, the phenological data comprised the leafout of vegetative buds in adult trees (Stations 1-3, Chinese National Ecology Science Data Center, www.cnern.org.cn). All temperature data were provided by the China National Meteorological Information Center (www.data.cma.cn).

| Station index | Phenological station |  |  | Meteorological station |  |  | Years | Tree species |
| --- | --- | --- | --- | --- | --- | --- | --- | --- |
| 1 | Changsha | 28°06' N | 113°00' E | Changsha | 28°07' N | 113°00' E | 2005-2008 | *Castanopsis*<br>*Phoebe*<br>*Pseudolarix* |
| 2 | Yinxian | 29°36' N | 121°12' E | Yinxian | 29°31' N | 121°20' E | 1984-1996 | *Pseudolarix* |
| 3 | Guilin | 25°13' N | 110°15' E | Guilin | 25°09' N | 110°06' E | 2005, 2006, 2008 | *Pseudolarix* |
| 4 | Guiyang | 26°34' N | 106°42' E | Guiyang | 26°21' N | 106°26' E | 2014-2016 | *Torreya* |

#### 2.4.4 Warming experiment

Lastly, the models were tested with independent bud burst data collected in a whole-tree chamber (WTC) warming experiment on the Zhejiang A&F University campus in autumn, winter and spring 2019 – 2020. In the experiment we applied a complete 3×3 random design with nine treatments combining three levels of both spring and winter warming: ambient temperature (A), ambient plus two degrees warming (A+2), and ambient plus four degrees warming (A+4). The winter and spring warming treatments were conducted from 10 November 2019 to 10 February 2020 and from 11 February 2020 until the observed bud burst, respectively. In each treatment we had eight replicated first-year seedlings of each of the four species. The seedlings had been raised like those used in the experiments carried out for model development (Zhang et al., 2021). Three WTCs (E-Lotus Technology Co., Beijing, China) were used in the experiment, one for each temperature level applied.

The nine temperature treatments were realized by moving seedlings from chamber to chamber. For each species, 24 seedlings were put in each of the three WTCs at the beginning of the experiment on 10 November 2019. On 11 February 2020, three sub-groups of eight seedlings were randomly sampled in each chamber. One sub-group remained in the same chamber and the other two were transferred, each into one of the WTCs representing the other two temperature levels. In this way, nine temperature treatments simulating the current climate and eight different cases of climatic warming were created. The air temperature records in each chamber were collected with iButtons (Model DS1912L, Embedded Data Systems Co., Ltd, KY, USA). Bud development was observed and the timing of bud burst was determined as they were in the experiments producing the experimental data for model development (Zhang et al., 2021).

### 2.5 Sensitivity analysis

In order to examine the potential role of temperatures below +5 °C in the rest break and the subsequent timing of bud burst, simulations of bud burst timing with long-term air temperature data as input were carried out with two models for each of the five material categories. In addition to the original model, we applied a modified model that assumed the rate of rest break to be at the maximum at any air temperature below +5 °C (see Figure 2 in the Results Section). We made use of the daily temperature records from Hangzhou, southeastern China (30°08′N, 120°06′E) for 1958 – 2018 by converting them into hourly temperatures by means of Eqn. (19).

### 2.6 Scenario simulations

The effects of climatic warming on the spring phenology were assessed by projecting the timing of bud burst for 2020 – 2099 by means of the NASA Earth Exchange Global Daily Downscaled Climate Projections (NASA NEX-GDDP) data set. This data set is comprised of downscaled climate scenarios for the globe that are derived from the General Circulation Model (GCM) runs conducted under the Coupled Model Intercomparison Project Phase 5 (CMIP5, Taylor, Ronald, & Meehl, 2011). We used daily minimum and maximum temperature records with a spatial resolution of 0.5×0.5° under two climatic scenarios, RCP 4.5 and RCP 8.5, (Meinshausen et al., 2011), which represented warming by 1.4 and 4.8 °C, respectively, for the period of 2020 – 2100 at our research site (Hangzhou). These data have been bias-corrected (Thrasher, Maurer, McKellar, & Duffy, 2012). The daily temperature records were converted into hourly temperatures by means of Eqn. (19). The predicted yearly times of leafout / flowering were plotted against time and, in order to quantify the effects of warming, a linear regression was fitted to the data.

### 2.7 The software used in the study

Most of the data analyses and model calculations were carried out with R statistical software v.4.0.2 (R Development Core Team, 2020). The sensitivity analyses were done with source codes written in Fortran with the Visual Studio 2010 software.

## 3 Results

### 3.1 Process-based models for the five material categories

#### Sub-model I

For *Torreya* trees, the experimental results show a clear decrease in the rate of rest break at temperatures upwards of +5 °C (Figure 2e; Table 4). A basically similar decrease was also found for *Pseudolarix* and *Torreya* seedlings, but because the decrease was less steep in these two material categories, their model formulation required extrapolation for temperatures above +15 °C (Section 2.3.2; Figure 2c,d; Table 4). In *Castanopsis* seedlings, a plateau response over the measured temperature range was found, and accordingly, a horizontal line was used in the curve fitting for the model formulation (Figure 2a; Table 4). A similar plateau response was also assumed for *Phoebe* seedlings, despite an outlier observed at +15 °C (Figure 2b; Table 4). For both *Castanopsis* and *Phoebe* seedlings, extrapolation was needed at temperatures above +15 °C (Figure 2a,b; Table 4). For temperatures below +5 °C, extrapolation was needed with all five material categories. For this low-temperature end, two optional responses were formulated for each material category (Figure 2; Table 4). The maximal rate of rest break was relatively low in *Castanopsis* seedlings (Figure 1a), high in *Phoebe* seedlings (Figure 2b) and *Torreya* trees (Figure 2e), and intermediate in *Pseudolarix* (Figure 2c) and *Torreya* (Figure 2d) seedlings.

**Table 4.** Equations of Sub-model I for the five material categories studied (seedlings of three species and both seedlings and twigs representing adult trees of *Torreya*). Sub-model I represents the air temperature response of the rate of rest break, R_r_ (function f_1_ in Eqn. 1 of the overall model; see Figure 2). In the modified model for each material category, the equations for all temperatures T < +5 °C are replaced by a constant value of R_r_ (dashed lines in Figure 2), either by using the constant value indicated by the shading (*Castanopsis* and *Phoebe* seedlings) or by calculating the constant value with the shaded equation with T = +5 °C (*Pseudolarix* and *Torreya* seedlings and *Torreya* trees).

| Material category | Sub-model I: $R_r$ (% h<sup>-1</sup>) = | |
| --- | --- | --- |
| *Castanopsis* seedlings | 0, | $T < -3.4$ °C |
| | $0.0077 T + 0.0263$, | $-3.4$ °C $\leq T < +5$ °C |
| | 0.0650, | $+5$ °C $\leq T < +15$ °C |
| | $-0.0130 T + 0.2600$, | $+15$ °C $\leq T < +20$ °C |
| | 0, | $T \geq +20$ °C |
| *Phoebe* seedlings | 0, | $T < -3.4$ °C |
| | $0.0183 T + 0.0623$, | $-3.4$ °C $\leq T < +5$ °C |
| | 0.1540, | $+5$ °C $\leq T < +15$ °C |
| | $-0.0308 T + 0.6160$, | $+15$ °C $\leq T < +20$ °C |
| | 0, | $T \geq +20$ °C |

| Material category | Sub-model I: $R_r$ (% $h^{-1}$) = | |
| --- | --- | --- |
| *Pseudolarix* seedlings | 0, | $T < -3.4\text{ }^{\circ}\text{C}$ |
| | $0.0094\text{ }T + 0.0319$, | $-3.4\text{ }^{\circ}\text{C} \leq T < +5\text{ }^{\circ}\text{C}$ |
| | $-0.0039\text{ }T + 0.0984$, | $+5\text{ }^{\circ}\text{C} \leq T < +15\text{ }^{\circ}\text{C}$ |
| | $-0.0080\text{ }T + 0.1596$, | $+15\text{ }^{\circ}\text{C} \leq T < +20\text{ }^{\circ}\text{C}$ |
| | 0, | $T \geq +20\text{ }^{\circ}\text{C}$ |
| *Torreya* seedlings | 0, | $T < -3.4\text{ }^{\circ}\text{C}$ |
| | $0.0150\text{ }T + 0.0509$, | $-3.4\text{ }^{\circ}\text{C} \leq T < +5\text{ }^{\circ}\text{C}$ |
| | $-0.0062\text{ }T + 0.1567$, | $+5\text{ }^{\circ}\text{C} \leq T < +15\text{ }^{\circ}\text{C}$ |
| | $-0.0127\text{ }T + 0.2548$, | $+15\text{ }^{\circ}\text{C} \leq T < +20\text{ }^{\circ}\text{C}$ |
| | 0, | $T \geq +20\text{ }^{\circ}\text{C}$ |
| *Torreya* trees | 0, | $T < -3.4\text{ }^{\circ}\text{C}$ |
| | $0.0185\text{ }T + 0.0627$, | $-3.4\text{ }^{\circ}\text{C} \leq T < +5\text{ }^{\circ}\text{C}$ |
| | $-0.0154\text{ }T + 0.2320$, | $+5\text{ }^{\circ}\text{C} \leq T < +15.1\text{ }^{\circ}\text{C}$ |
| | 0, | $T \geq +15.1\text{ }^{\circ}\text{C}$ |

#### Sub-model II

For the air temperature response of the potential rate of ontogenetic development, R_o,pot_, the sigmoidal equation fitted to the results for all five material categories quite well in general (Figure 3; Table 5). At high temperatures, however, there was a great deal of variation in the R_o,pot_ values among the *Castanopsis* (Figure 3a) and *Pseudolarix* (Figure 3c) seedlings. In the estimated air temperature response of R_o,pot_, there were marked differences among the material categories (Figure S3). For *Castanopsis*, *Phoebe*, and *Torreya* seedlings, the lower temperature threshold for ontogenetic development was estimated to be approximately +10 °C, whereas for *Torreya* trees and *Pseudolarix* seedlings, the rate of ontogenetic development was estimated to be considerable at temperatures below +10 °C also (Figure 3, Figure S3). For *Torreya* seedlings, the curve was estimated to level off at +20 °C already, and their rate of development was also estimated to be generally lower than that of the other material categories (Figure 3, Figure S3). For *Castanopsis, Phoebe*, and *Pseudolarix* seedlings, the curve levelled off near +30 °C, whereas for *Torreya* trees, no levelling off was found (Figure 3, Figure S3).

**Table 5.** Parameter values of Sub-model II for the five material categories studied (seedlings of three species and both seedlings and twigs representing adult trees of *Torreya*). Sub-model II represents the sigmoidal air temperature response of the potential rate of ontogenetic development, R_o,pot_ (function f_2_ in Eqn. 4 of the overall model; see Figure 3). The potential rate of ontogenetic development indicates the rate after rest completion, when there is no longer any reduction of the rate caused by the rest status of the bud. The parameters a_2_, b_2_, and c_2_ indicate the upper asymptote (maximal rate), steepness, and inflexion point of the sigmoidal curve, respectively.

| Material category | $a_2$ (% h<sup>-1</sup>) | $b_2$ (°C<sup>-1</sup>) | $c_2$ (°C) | $R^2$ |
| --- | --- | --- | --- | --- |
| *Castanopsis* seedlings | 0.3984 | -2.9868 | 20.0907 | 0.8518 |
| *Phoebe* seedlings | 0.3451 | -3.6026 | 20.5672 | 0.9041 |
| *Pseudolarix* seedlings | 0.3008 | -3.6984 | 14.4639 | 0.6125 |
| *Torreya* seedlings | 0.1130 | -1.8091 | 17.2439 | 0.9401 |
| *Torreya* trees | 0.2709 | -8.8756 | 21.3366 | 0.8742 |

#### Sub-model III

Considerable differences were found among the material categories in the estimated dependence of ontogenetic competence, C_o_, on the state of rest break, S_r_ (Figure 4; Table 6). In broad terms, three patterns are seen in the results. First, in *Torreya* trees, C_o_ stays at zero until S_r_ reaches about 68 %, after which C_o_ abruptly attains its near-maximal value (Figure 4e). Second, similarly to *Torreya* trees, in *Pseudolarix* (Figure 4c) and *Torreya* (Figure 4d) seedlings, C_o_ stays at zero with low values of S_r_, but in contrast to *Torreya* trees, C_o_ gets positive values as soon as S_r_ reaches about 25 %. After that, C_o_ increases with increasing values of S_r_ in these two material categories. Third, in *Castanopsis* (Figure 4a) and *Phoebe* (Figure 4b) seedlings, C_o_ has a positive value even at S_r_ = 0 %, indicating that in these two species leafout can take place even without chilling. Considerable differences among the five material categories were also found in the slope of the dependence of C_o_ on S_r_ (Figure 4; Table 6).

**Table 6.** Parameter values of Sub-model III (Eqn. 18) for the five material categories studied (seedlings of three species and both seedlings and twigs representing adult trees of *Torreya*). Sub-model III mediates the effect of the rest status on the rate of ontogenetic development towards bud burst (Figure 1) by formulating the dependence of ontogenetic development, C_o_, on the state of rest break, S_r_ (function f_3_ in Eqn. 5 of the overall model; see Figure 4).

| Material category | Parameter |  |  |
| --- | --- | --- | --- |
| | $a_3$ | $b_3$ | $S_{r,min}$ |
| *Castanopsis* seedlings | 0.0076 | 0.2410 | 0 |
| *Phoebe* seedlings | 0.0037 | 0.6334 | 0 |
| *Pseudolarix* seedlings | 0.0082 | 0.1765 | 20.16 |
| *Torreya* seedlings | 0.0044 | 0.5593 | 24.85 |
| *Torreya* trees | 0.0050 | 0.4974 | 67.29 |

### 3.2 Tests of the models with independent bud burst data

#### 3.2.1 Predicted dormancy dynamics and a test with short-term phenology records

The dormancy dynamics implied by the ecophysiological phenomena addressed by the three sub-models are relatively complicated, giving rise to predictions that often cannot be discovered, not even in qualitative terms, without computer simulations. These dynamics are illustrated in Figure 5 for the five material categories addressed in the present study.

**FIGURE 5.**
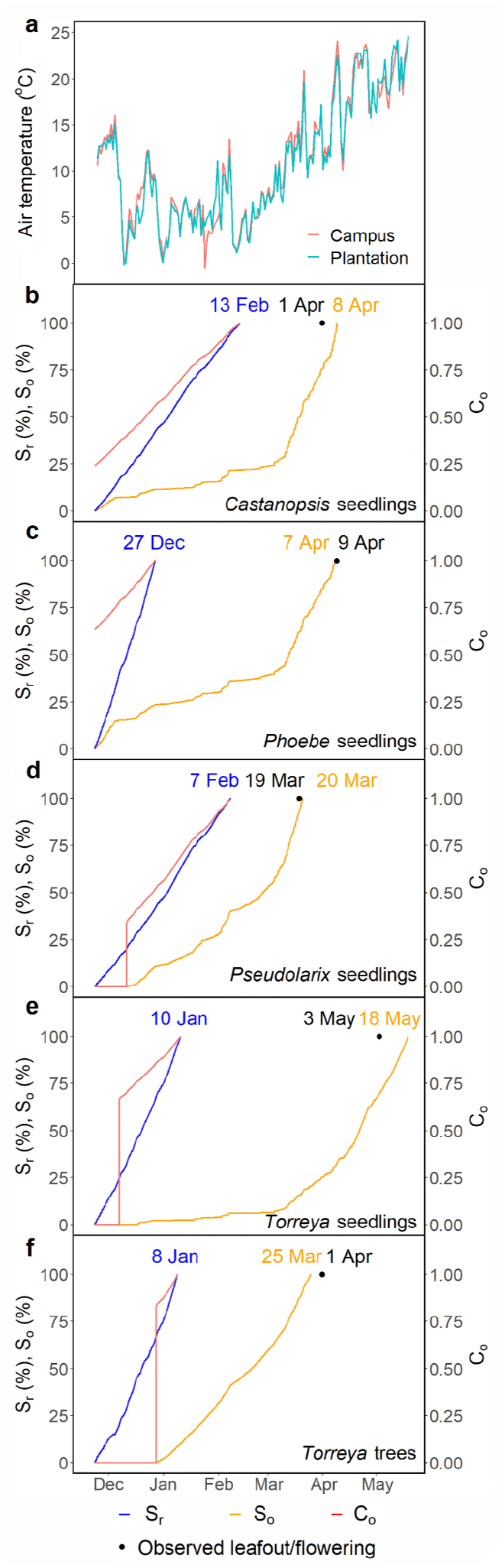
(a) Daily mean temperatures for the campus seedling collection (for seedlings of the four species in panels b-e) and for the *Torreya* tree plantation (for *Torreya* trees in panel f). (b-f) Predicted dormancy dynamics for autumn 2018 and spring 2019 of the five material categories addressed in the study. The vertical axis on the left represents the state of rest break, S_r_, (blue line) and the state of ontogenetic development, So, (orange line). Rest completion is predicted to occur when S_r_ = 100 % (date marked in a blue font) and leafout/flowering when So = 100 % (date marked in an orange font). The observed leafout/flowering is marked with a circle and the date in a black font. The vertical axis on the right represents ontogenetic competence, C_o_, (red line). By definition, C_o_ = 1 at rest completion, when S_r_ = 100 %. Otherwise the relationship between C_o_ and S_r_ varies among the five material categories as shown in Figure 4

For all five material categories, rest break was predicted to start at the beginning of the simulation on November 23, as indicated by the increasing values of the state of rest break, S_r_ (Figure 5b-f). This was because at the time, the air temperature was mainly fluctuating in the rest-breaking range already (Figure 5a; compare with Figure 2). Rest break was the fastest in *Phoebe* seedlings, where rest completion was predicted to occur on 27 December (Figure 5c). This is because *Phoebe* seedlings retain their high rate of rest break over a broad temperature range (Figure 2b). In the other material categories, rest completion varied from 8 January in *Torreya* trees (Figure 5f) to 13 February in *Castanopsis* seedlings (Figure 5b).

Besides the predicted S_r_, the rate of ontogenetic development is also affected by two other phenomena. First, there are differences among the material categories in the dependence of ontogenetic competence, C_o_, on S_r_ (Sub-model III; Figure 4); thus *Torreya* seedlings, for instance, started their ontogenetic development earlier (with S_r_ = 24.85 %, Figure 4d) than *Torreya* trees (with S_r_ = 67.29 %, Figure 4e), even though no major differences in rest break (development of S_r_) were seen between them (Figure 5e,f). Second, even when two material categories have the same C_o_, there may be differences in their rate of ontogenetic development (Sub-model II, Figure 3, Figure S3). *Torreya* seedlings, for instance, show an exceptionally low rate of ontogenetic development (Figure S3), and accordingly, despite their earlier start of ontogenetic development, the *Torreya* seedlings showed predicted bud burst later than the *Torreya* trees showed predicted flowering (Figure 5e,f).

In predicting the timing of spring phenology, the highest accuracy was reached with *Pseudolarix* seedlings, where the prediction was one day late (Figure 5d). *Pseudolarix* seedlings were followed by *Phoebe* seedlings (two days early, Figure 5c), *Castanopsis* seedlings (seven days late, Figure 5b), *Torreya* trees (seven days early, Figure 5f), and *Torreya* seedlings (fifteen days late, Figure 5e).

#### 3.2.2 Independent tests with phenology records and results of the warming experiment

The models developed for *Torreya* provided relatively accurate predictions for both the timing of flowering in the observational phenological records and the timing of seedling leafout found in the warming WTC experiment of the present study (Figure 6d). For *Torreya*, the predictions were also unbiased.

**FIGURE 6.**
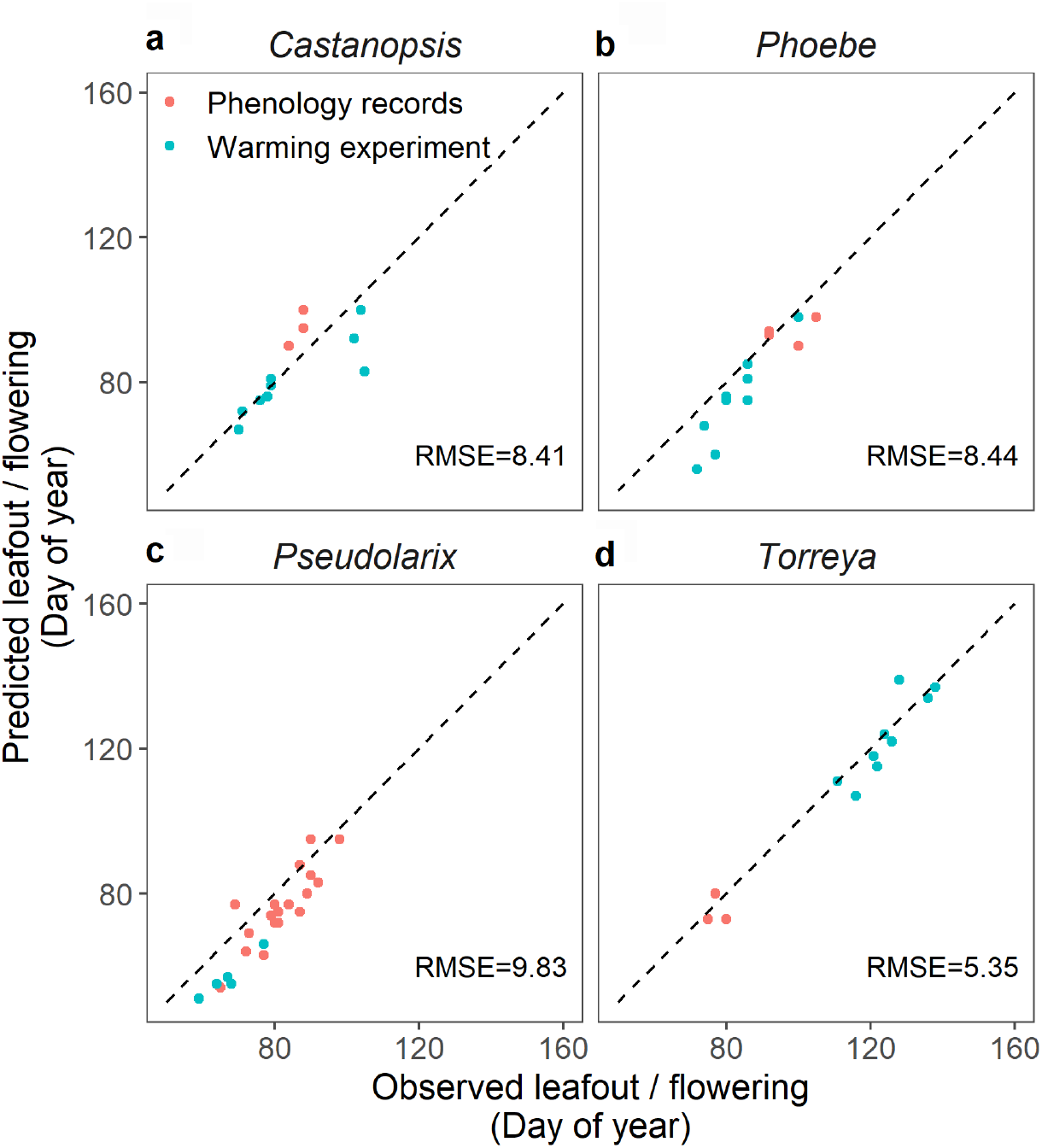
Independent tests of the process-based spring phenology models developed for the four subtropical tree species by using both phenological records and results from our warming whole-tree chamber (WTC) experiment. The phenological records represent adult trees (flowering in *Torreya*, leafout in the other three species), whereas the results from the WTC experiment represent leafout in the seedlings of all four species. The predictions for the phenological records of *Castanopsis, Phoebe*, and *Pseudolarix* trees were calculated by means of the models developed for seedlings because no model for adult trees had been developed for these species in the present study

For the other three species, no model for adult trees was developed in the present study, so that for predicting the timing of leafout in adult trees documented in the phenological records, the seedling leafout model was used. In comparison with *Torreya*, the accuracy reached with the other three species was lower (Figure 6a,b,c). Additionally, a bias was introduced into the model predictions for *Phoebe* (Figure 6b) and *Pseudolarix* (Figure 6c) as the model predictions were systematically earlier than the observations. No clear accuracy difference between the predictions of leafout for adult trees and leafout for seedlings (phenology records vs. warming experiment, Figure 6a,b,c) was found in any of these three species.

### 3.3 Sensitivity analysis

In the sensitivity analysis of the role that temperatures below +5 °C in rest break may have on the timing of the subsequent bud burst, considerable differences were found among the five material categories (Table 7). In *Torreya*, *Phoebe*, and *Castanopsis* seedlings, a negligible sensitivity to the difference represented by the two alternative models used as Sub-model I was found. The percentage of the years with a difference in the timing of bud burst found between the two alternative sub-models in *Torreya, Phoebe*, and *Castanopsis* seedlings was 1.6 %, 6.6 %, and 19.7 % respectively, but in all years where there was a difference, the modified model predicted bud burst to occur only one day earlier than the original model (Table 7). *Pseudolarix* seedlings displayed an intermediate sensitivity, with a difference between the predictions of the alternative models appearing in 77 % of the years. Between the results obtained with the two alternative sub-models, the average difference was 1.9 days and the maximum difference 7 days. *Torreya* trees showed the highest sensitivity, with a difference between the two simulations in almost all years, with the average and maximal differences of 2.5 and 17 days respectively, between the results obtained with the two alternative sub-models (Table 7).

**Table 7.** Results of the sensitivity analysis of the potential rest-breaking effects of temperatures below +5 °C on the timing of bud burst in the subsequent spring. The timing of bud burst in Hangzhou, southeastern China, was simulated for 61 years by using two alternative models for the temperature response of rest break. In the original model, no rest was assumed to take place at temperatures below −3.4 °C, whereas in the modified model, rest break was assumed to progress at its maximal rate at all temperatures below +5 °C (Figure 2; Table 4). The results in the table are for the years when there was a difference in the prediction between the two models. The positive values of the difference indicate that the modified model predicted an earlier bud burst than the original model.

| Material category | Percentage of years<br>with a difference in<br>the prediction | Mean difference<br>(days) | Maximal difference<br>(days) |
| --- | --- | --- | --- |
| *Castanopsis* seedlings | 19.7 | 1 | 1 |
| *Phoebe* seedlings | 6.6 | 1 | 1 |
| *Pseudolarix* seedlings | 77.0 | 1.9 | 7 |
| *Torreya* seedlings | 1.6 | 1 | 1 |
| *Torreya* trees | 96.7 | 2.5 | 17 |

### 3.4 Scenario analyses

The timing of spring phenology was projected to advance in all scenario simulations that addressed the effects of climatic warming in 2020 – 2100 (Figure 7, Figure S4). Among the seedlings of the four species, the advances in the rates of leafout varied between 0.4 and 0.8 days per decade for RCP4.5 and between 2.8 and 3.5 days per decade for RCP8.5. The projected advances in seedling leafout per one °C warming varied between 4.7 and 5.9 days per °C, with no major differences between the two climatic scenarios (Figure 7a,b,c,d). For *Torreya* flowering, the results were different from those for leafout in the seedlings: first, a relatively minor advance of 0.4 and 0.9 days per decade was projected for RCP4.5 and RCP8.5, respectively. Second, there was a major difference between the advances per °C warming projected for RCP4.5 (2.3 days per °C) and for RCP8.5 (0.9 days per °C) (Figure 7e).

**FIGURE 7.**
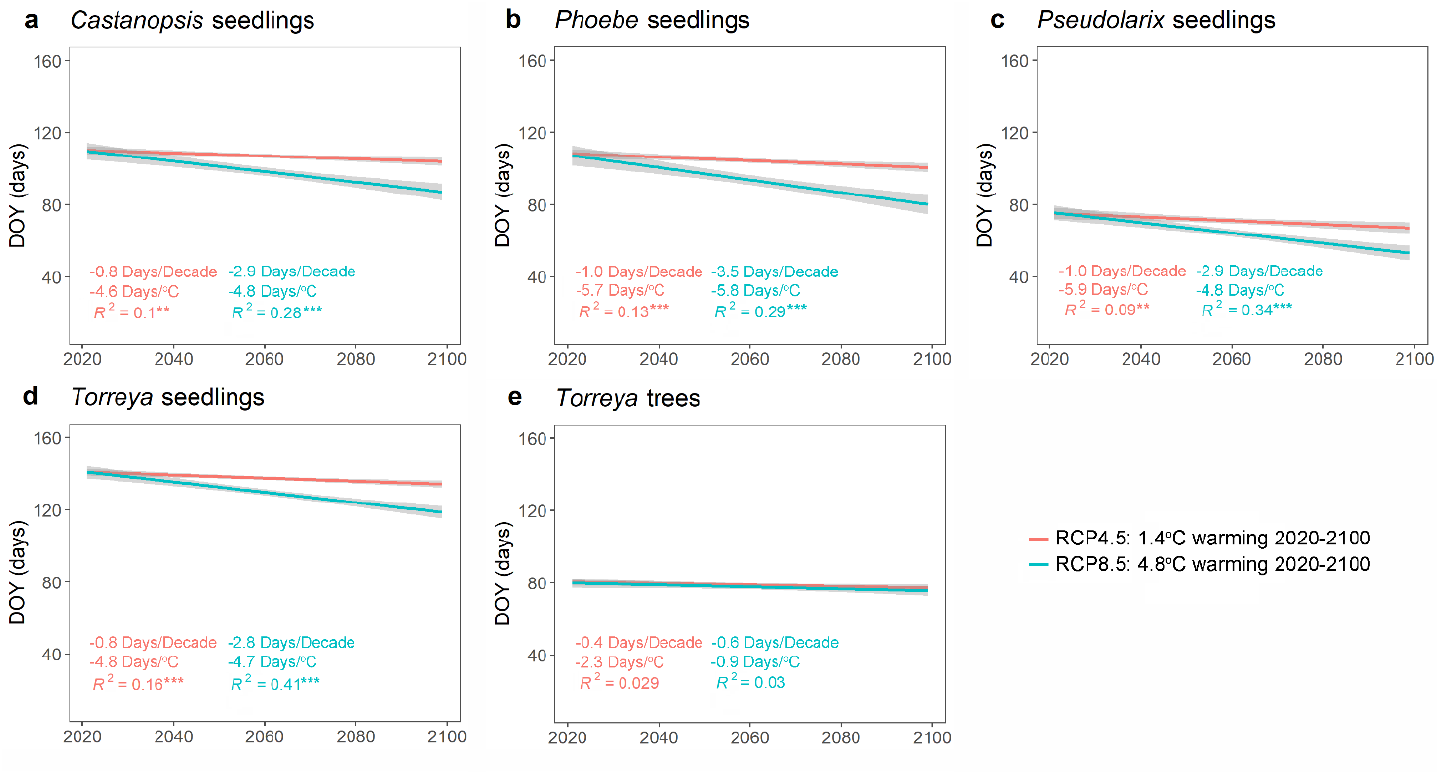
7 Projected timing of spring phenology (flowering for *Torreya* trees, leafout for the seedlings of the four species) for 2020 – 2100 under the climatic scenarios RCP4.5 and RCP8.5. The projections were calculated by means of the process-based models developed in the present study. The trends are presented here with direct lines fitted to the results; for the original yearly results, see Figure S4

## 4 DISCUSSION

### 4.1 Biological realism of the models

We introduced novel process-based models of spring phenology for four subtropical tree species. The main emphasis in our modelling was on the biological realism of the models (Levins, 1966, 1968; Charrier, Ngao, Saudreau, & Améglio, 2015; Hänninen, 2016), which is why we took an experimental approach explicitly addressing the ecophysiological phenomena modelled. This approach, erred to as *the ecophysiological approach* by Hänninen (2016), has been taken with boreal and temperate trees only rarely (Sarvas, 1972, 1974; Campbell & Sugano, 1975,1979; Caffarra, Donnelly, & Chuine, 2011) and, to our knowledge, with subtropical trees never before. The obvious reason for the shortage of earlier studies is the large amount of labour required by the experiments. The modular model structure adopted in the present study (Figure 1) facilitates the designing of particular experiments with each one addressing a specific ecophysiological phenomenon that affects the timing of the spring phenological event. The modular model structure was introduced as early as 30 years ago, but as far as we know, our study is the first one where it was used to its full potential, basing all three submodels on specific experimental data for the species addressed.

In the process-based modelling of tree spring phenology, an approach partially based on previous experimental findings is sometimes taken. In this approach, referred to as *the intermediate approach* by Hänninen (2016), rest break is simulated by accumulating chilling units and ontogenetic development by accumulating forcing units, with both based on temperature responses previously determined experimentally for some other species than the ones being modelled. In that approach, only the chilling and forcing requirements are determined on the basis of species-specific data (Häkkinen, Linkosalo, & Hari, 1998; Luedeling, Zhang, McGranahan, & Leslie, 2009; Xu, Dai, Ge, Wang, & Tao, 2020). With reference to the temperature responses determined in the present study (Figures 2, 3), this would mean that the shape of the temperature response curve is fixed *a priori* but the level is estimated from new species-specific data: the higher the chilling/forcing requirement, the lower the level of the corresponding curve (Hänninen, 2016). In a similar approach often taken, not even the shapes of the temperature response curves have any experimental basis. For instance, the threshold chilling model, which assumes that one chilling unit is accumulated during each hour when the air temperature is below a given threshold, is used quite frequently (Weinberger, 1950; Cannell & Smith, 1983), but as far as we know, no explicit experimental data has been published to document a response of that type.

Our explicit experimental results show that there are considerable differences among the examined four subtropical tree species in their air temperature responses of the rate of rest break (Figure 2) and ontogenetic development (Figure 3). Similar differences were found between the two life stages of *Torreya* addressed (Figures 2e,f, 3e,f). Even bigger differences were found between our results for the subtropical trees and those found earlier for boreal and temperate trees. Most importantly, we found that temperatures of up to +15 °C are effective in the rest break of many subtropical tree species. In fact, with the exception of adult *Torreya* trees, we were not even able to determine the upper limit of the rest-breaking temperature range for the subtropical tree species we studied (Zhang et al., 2021). In most of the previous experimental studies of more northern trees, the upper threshold was found to be +10 to +12 °C or lower (Sarvas, 1974; Heide & Prestrud, 2005, however, see Erez & Couvillon, 1987). Similarly, we found in the present study that temperatures between 0 and +10 °C, which have been found to promote ontogenetic development towards bud burst in boreal and temperate trees (Sarvas, 1972), are ineffective for many subtropical trees. All of these comparisons indicate that there is no chilling unit for rest break and no forcing unit for ontogenetic development based on response curves fixed *a priori* that could be universally valid for all species. Rather, the modelling needs to be based on species-specific or possibly even on provenance-specific data (Hänninen, 2016).

In seeking to attain species-specific models, an approach referred to as *the phenological approach* by Hänninen (2016) is often taken. In this approach, process-based tree phenology models are developed with the technique of inverse modelling, which means that rather than basing the models and their parameter values on experimental studies explicitly designed to examine the processes involved, the models and their parameter values are determined by fitting the models statistically to observational long-term phenological records (Kramer, 1994; Chuine, Cour, & Rousseau, 1998, Basler, 2016). The approach has become popular as it requires no experimental work and as long-term phenological records are nowadays readily available for several tree species and geographical locations. Accordingly, when developing the first process-based model for a subtropical tree species, Chen et al. (2017), too, took the observational approach by using inverse modelling. However, as already shown by the pioneering study of Hunter and Lechowicz (1992), inverse modelling involves an exceptionally high degree of uncertainty. It often produces biologically unrealistic models, such as the ones suggesting that temperatures above 100 °C belong to the chilling temperature range (Kramer, 1994; for a recent discussion of the problems related to inverse modelling, see Hänninen et al., 2019). Unfortunately, the study of Chen et al. (2017), too, was undermined by unrealistic temperature responses of chilling accumulation (Figure S5).

### 4.2 Further model development

Though our experimental approach explicitly addressed the processes being modelled, the models developed still require further work for better biological realism. Several additional aspects need to be addressed in further work of process-based modelling for subtropical trees. The most obvious one is seen in Figure 2, which shows that in the sub-model for the air temperature response of the rate of rest break, extrapolation was generally needed at both the low and the high end of the temperature spectrum. However, when experiments are carried out to add the missing data points, various methodological problems are met. Temperatures above +15 °C generally cause ontogenetic development towards bud burst, thus lowering the value of DBB and confusing the determination of the chilling duration required for rest completion. This problem can be addressed with computational techniques, such as simulated annealing (Chuine et al., 1998; Chuine, 2000; Lundell, Hänninen, Saarinen, Åström & Zhang, 2020). At the lower end of the temperature scale, problems may be met in chilling with freezing temperatures because subtropical tree species do not necessarily survive in continuous freezing conditions (Zhang et al., 2021). The sensitivity analyses of the present study also suggest that temperatures below +5 °C occur so rarely at the subtropical research site that their effect on rest break may be quite limited with most species. Still, at least the temperatures between zero and +5 °C need to be studied for their rest-breaking effects.

In addition, the effects of fluctuating temperatures (Fishman, Erez & Couvillon, 1987a,b) and of photoperiod on rest break (Zhang et al., 2020) need to be introduced into the process-based modelling for subtropical trees. Furthermore, the post-rest theory introduced by Vegis (1964) and recently formulated for boreal field layer plants by Lundell et al. (2020) should be examined with reference to subtropical trees, too. The theory suggests that the temperature range promoting ontogenetic development changes during dormancy. That change provides a potential explanation for the bias of the model prediction observed in the present study with two species in the independent test (Figure 6; Hänninen, 2016). The generic concept of ontogenetic competence (Figure 1) used in the present study was also used by Lundell et al. (2020), showing that the concept can be applied in a broad range of studies where restrictions caused by the rest status on the ontogenetic development need to be addressed.

Similarly to most studies of process-based tree phenology modelling, an indirect black-box modelling approach was taken in the present study, too, in that rather than considering the processes of rest break and ontogenetic development as such, only their final outcome (leafout, flowering) was observed in the experiments. For ontogenetic development, the processes could be addressed by microscopic observations of the anatomical development of the bud (Sarvas 1972, 1974;

Sutinen, Partanen, Viherä-Aarnio, & Häkkinen, 2012; Viherä-Aarnio, Sutinen, Partanen, & Häkkinen, 2014). This is because the models predict not only the occurrence of leafout/flowering when So = 100 % but also the occurrence of a given anatomical stage of development when So attains a constant critical value (< 100 %) specific to the stage of development. Despite our improved understanding of the physiological mechanisms of rest and rest break (Rohde & Bhalerao, 2007; Cooke, Eriksson, & Junttila, 2012; Tylewicz et al., 2018), no corresponding markers have been available for similar testing of the models of rest break in regard to developmental events corresponding to S_r_ values below 100 %. Chuine et al. (2016), however, suggested the use of the floral primordia fresh weight as such a marker for *Prunus armeniaca*. It would seem useful to examine that marker’s suitability for other species as well.

### 4.3 Projected effects of climatic warming

In our scenario analyses, an advancing of the spring phenology was projected for all five material categories addressed (Figure 7). For the seedlings of the four species, the projected advancing per one °C warming varied between 4.7 and 5.9 days per °C, which is in line with the corresponding projections obtained in earlier studies for temperate tree species (Kramer, 1994; Murray et al., 1994; Wang et al., 2020). For *Torreya* flowering, however, a smaller advancing was projected, and the advancing per one °C warming was much less for RCP8.5 than for RCP4.5 (Figure 7). In accordance with our hypothesis, the deviating results projected for *Torreya* flowering are explained by the peculiar aspects of the dormancy processes in *Torreya* flower buds, which are different from the corresponding processes in the vegetative buds of the seedlings of the four species studied.

First, rest break in *Torreya* trees proceeds relatively slowly at +10 °C, and at temperatures above +15 °C, no rest break is seen (Figure 2d). In the future, then, an increasing part of the winter will be too warm for rest break in *Torreya* flower buds, setting *Torreya* trees apart from the seedlings of the four species, especially those of *Castanopsis* and *Phoebe*. Second, rest break in *Torreya* trees needs to progress into an exceptionally late phase before any ontogenetic development is seen (Figure 4). This amplifies the effect of reduced chilling on its springtime phenology. Third, in comparison with the seedlings of the four species, the rate of ontogenetic development in *Torreya* trees shows a smaller increase in the temperature range of +10 to +20 °C (Figure S3), which is critical under climatic warming.

The deviating response of *Torreya* tree flowering to warming shows once again that process-based models should be based on specific responses determined by means of experiments explicitly designed to determine the responses of different species, also in different life stages of the given species. No modelling based on temperature responses fixed *a priori* could have revealed the deviating response to warming found for *Torreya* flowering.

### 4.4 Concluding remarks

We developed process-based models of spring phenology for four subtropical tree species. The development of the models was based entirely on specific experiments designed to address the processes, one at a time, that affect the spring phenology. Mostly, reasonable accuracy was attained in testing the models with independent data, both observational and experimental. In the application of the models to scenario simulations addressing the effects of climatic warming, an advancing of spring phenology was projected. Much less advancing was projected for flowering in *Torreya* than for leafout in the seedlings of the four species studied, and this was particularly the case under pronounced climatic warming. This difference was explained with reference to the peculiar dormancy processes typical for the flower buds of *Torreya* trees that were revealed in the experimentally based modelling. This shows the power of the experimental approach and, at the same time, calls for further experimental studies, our study being the first of its kind with subtropical trees.

## ACKNOWLEDGEMENTS

We thank Pekka Hirvonen (www.toisinsanoen.eu) for revising the language of the manuscript.

## AUTHOR CONTRIBUTION

R.Z., J.W. and H.H. designed the study. J.L., F.W. and R.Z. carried out the experiments. J.L., R.Z. and H.H. analyzed the data and carried out the modelling. R.Z., H.H. and J.L. wrote the study with inputs from J.W.

## DATA AVAILABILITY STATEMENT

The data that support the findings of this study are available from the corresponding author upon reasonable request.

