## Supplemental Information for "Effects of climatic warming on spring phenology in subtropical trees: process-based modelling with experiments designed for model development"

### Supplementary material

Rui Zhang, Jianhong Lin, Fucheng Wang, Heikki Hänninen & Jiasheng Wu

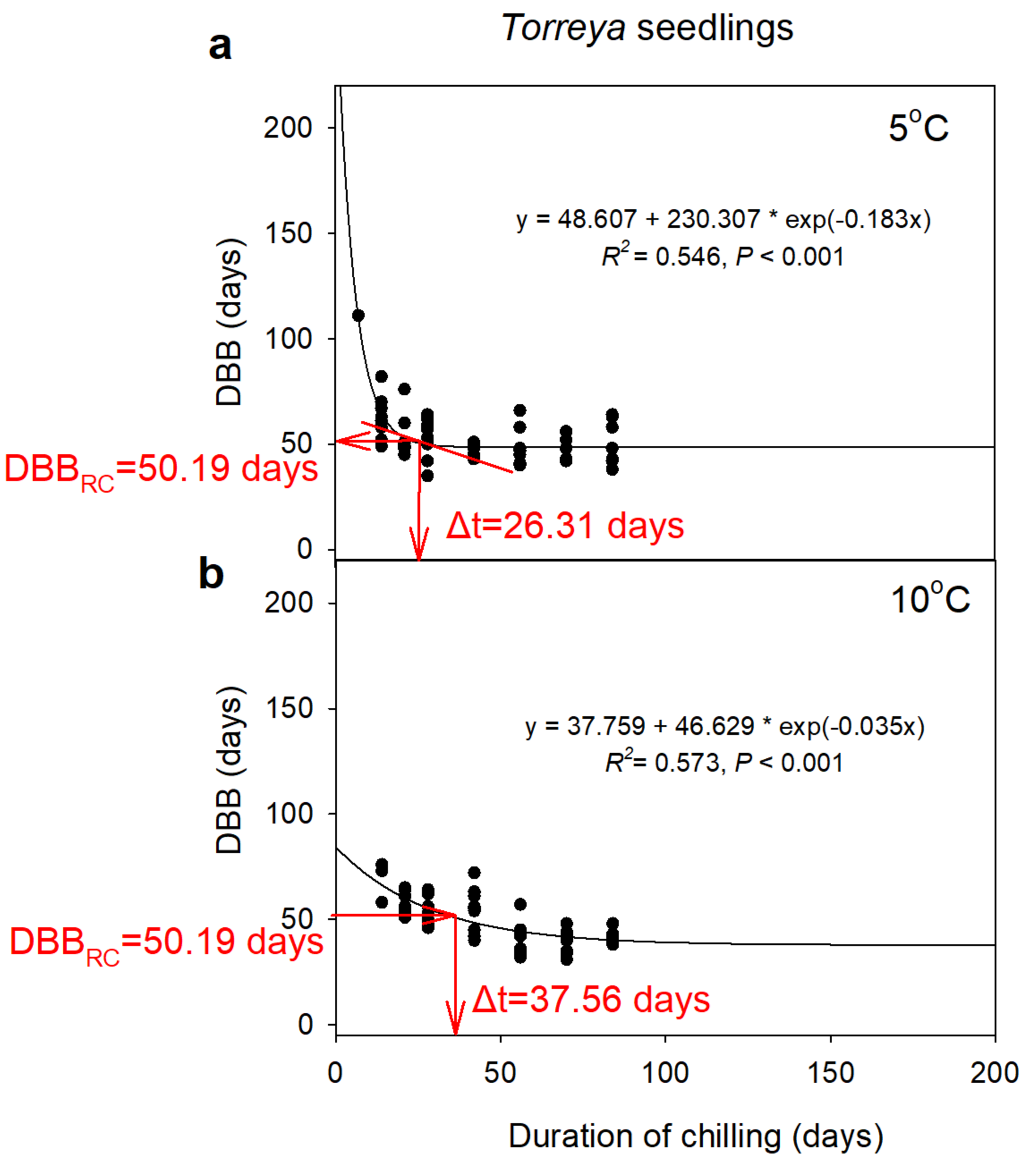


**FIGURE S1** The principle of determining the time in chilling conditions required for rest completion, Δt. (a) For chilling at +5 °C, Δt was determined as the duration of chilling corresponding to the slope of -0.3 (days/days) of the tangent of the DBB curve. The corresponding DBB value is denoted as $DBB_{RC}$, with RC referring to rest completion. (b) For chilling at +10 °C and at +15 °C, Δt was determined as the duration of chilling corresponding to $DBB_{RC}$. Data from Zhang et al. (2021)

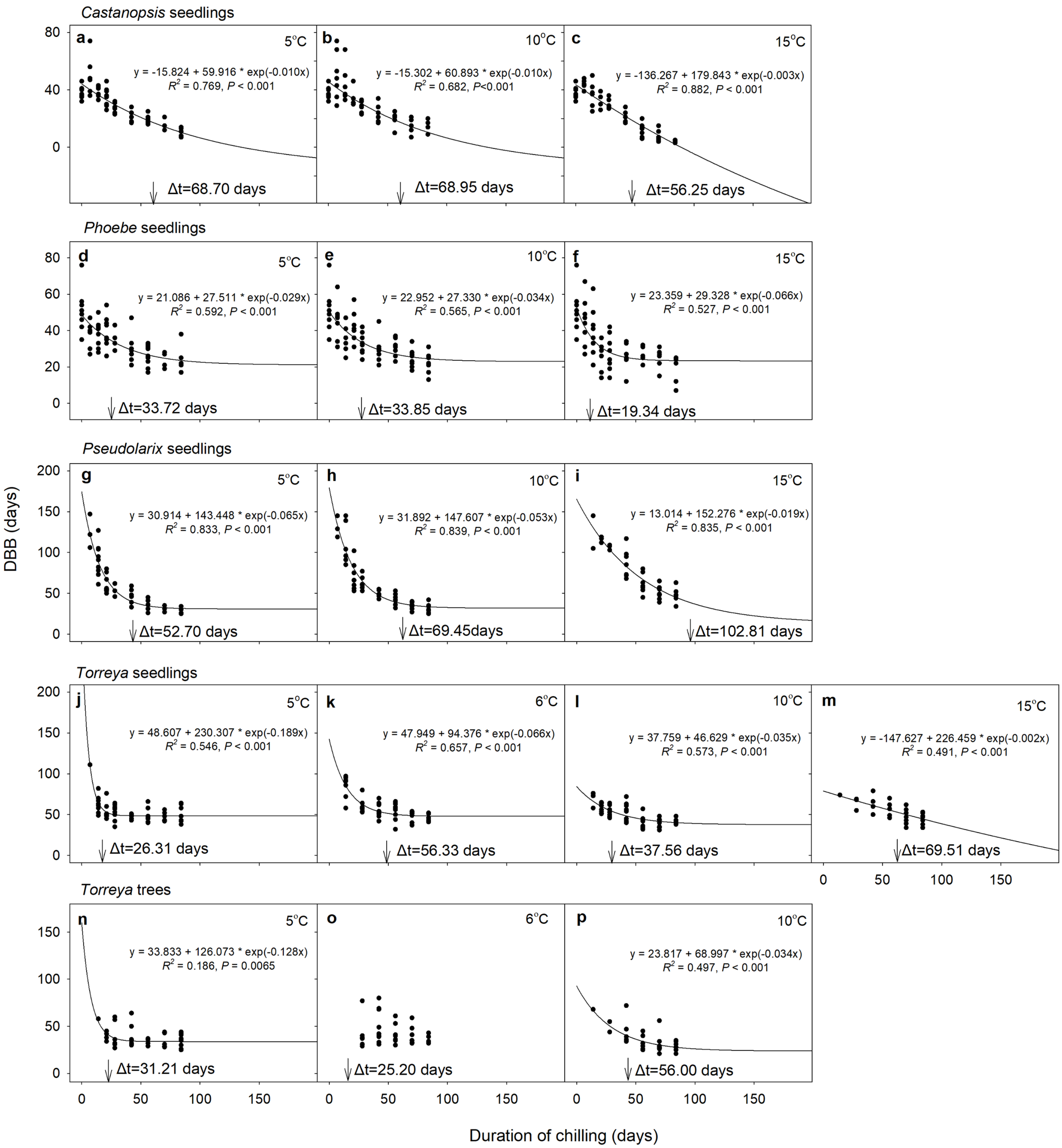


**FIGURE S2** Determination of the time, Δt, required for rest completion in chilling conditions for the five material categories in the various chilling conditions. In addition to the criterion based on the DBB curves (Figure S1), it was also required for Δt that BB% ≥ 80 % (BB% = bud burst percentage). That BB% criterion determined the Δt in two cases only. First, with *Torreya* tree chilling at +6 °C, no exponential decline of the DBB values was observed, so that the BB% criterion determined the value of Δt without any further consideration of DBB (panel o). Second, with *Torreya* tree chilling at +10 °C, BB% was below 80 % at the chilling duration where the criterion set for DBB (Figure S1b) was met, so that the value of Δt was determined by the BB% criterion (panel p). In all other cases the criterion based on DBB implied a higher value of Δt than the one based on BB%. This is why the BB% curves reported by Zhang et al. (2021) are not reported here. Please note the differences in the scaling of the vertical axes among the material categories. Data from Zhang et al. (2021)

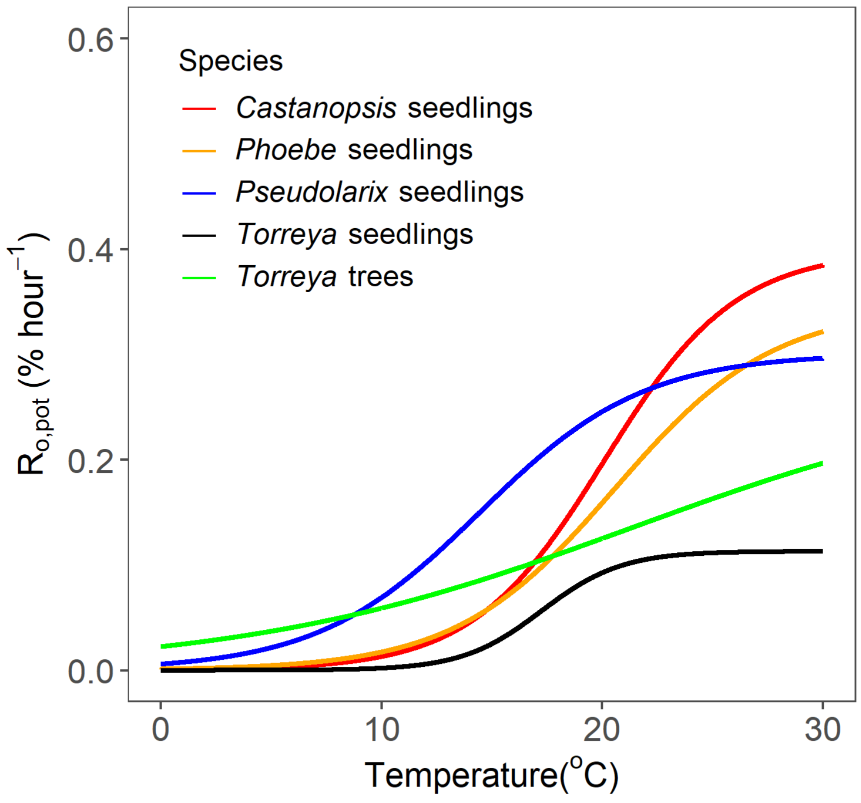


**FIGURE S3** A comparison of Sub-model II, formulated in the present study on the basis of experimental data, among the five material categories. $R_{o,pot}$ = potential rate of ontogenetic development

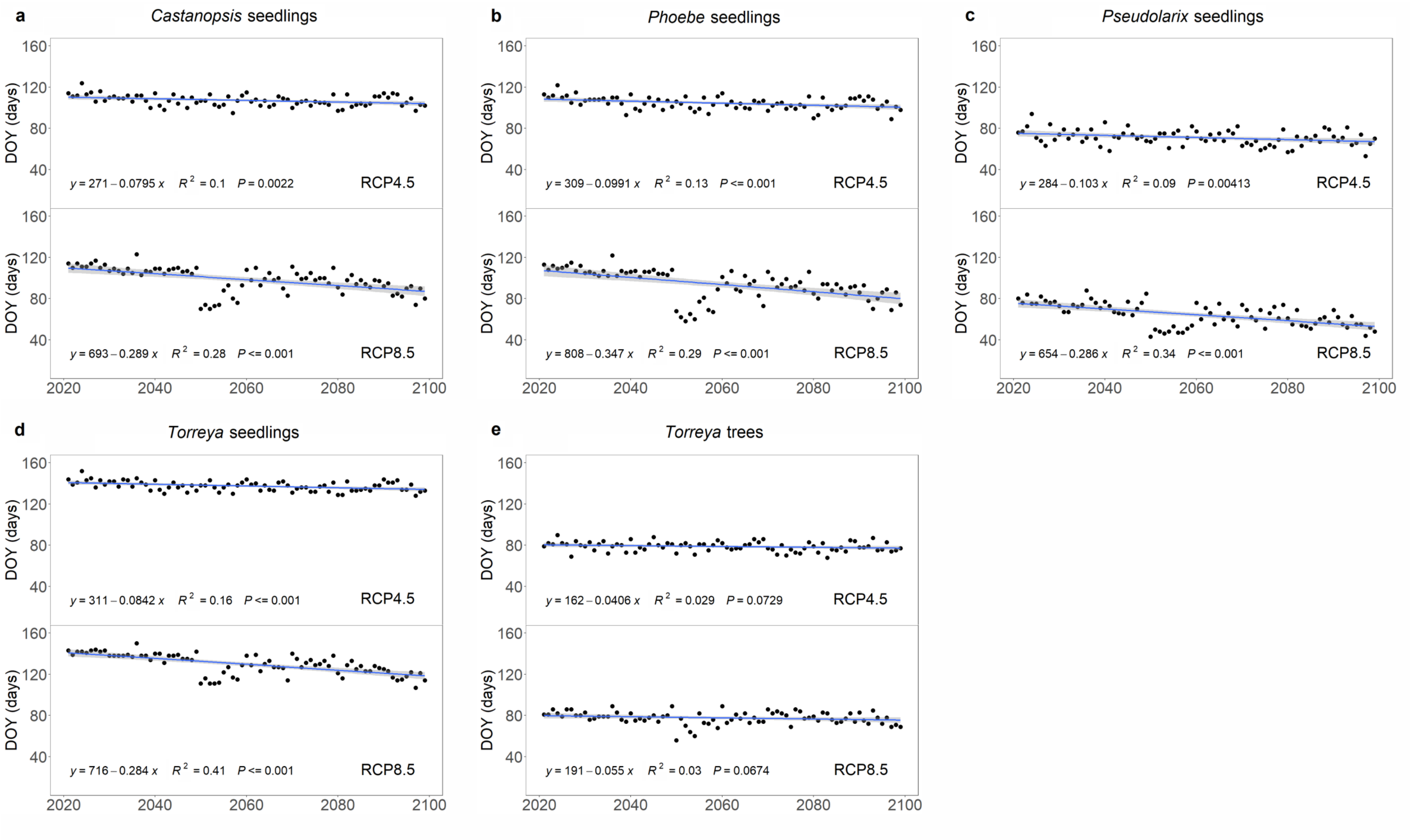


**FIGURE S4** Projected timing of spring phenology in the five material categories for 2020 – 2100 under the climatic scenarios RCP4.5 and RCP8.5. The Day of Year (DOY) (a-d) of leafout in seedlings of the four species and (e) of female flowering in *Torreya* trees

**FIGURE S5** (next page) Examples of the temperature responses estimated by Chen et al. (2017) for the rate of rest break (rate of chilling accumulation) in *Melia azedarach* growing in subtropical and northern tropical China. Chen et al. (2017) used the UniChill model (Chuine, 2000):

$$R_c(x_t) = \frac{1}{1 + e^{a(x_t - c)^2 + b(x_t - c)}}$$

where $R_c(X_t)$ = the rate of chilling accumulation at temperature $X_t$. Chen et al. (2017) fitted the equation to observational phenological data by estimating the values of parameters a, b, and c. The equation was fitted separately to the data representing various phenological events and phenological observation stations, which are both indicated in the panels. The parameter values were tabulated, but no curves were presented by Chen et al. (2017). Here we present a few examples of the curves implied by the tabulated parameter values by classifying the responses into four response types. (a-d) Type 1: no temperature response, chilling accumulation at a constant rate regardless of the prevailing temperature. (e-h) Type 2: high temperatures accumulate chilling. (i-l) Type 3: intermediate temperatures accumulate chilling. (m-p) Type 4: low temperatures accumulate chilling. Types 1 and 2 are not consistent with the concept of chilling requirement. Furthermore, several examples of the more realistic types 3 and 4 also show unrealistic details. The effective chilling range is often so wide in these response types that the responses become almost similar to the Type 1 responses (panels m and p). At the other extreme is the first leaf unfolding at station 23 in that its UniChill model suggests chilling accumulation in the temperature range of 3 to 5.5 $^{o}$C only (panel j). Obviously, such a narrow range cannot represent the real physiological response of the trees. In some cases a different response was found for the same phenological event at two stations located in the same climatic zone (panels c and g). In some other cases the response changed from the first leaf unfolding (panel f) to 50 % leaf unfolding (panel c) at the same station. In conclusion, these examples show that the inverse modelling used by Chen et al. (2017) unfortunately produced inconstant and biologically unrealistic temperature responses of the rate of rest break (rate of chilling accumulation)

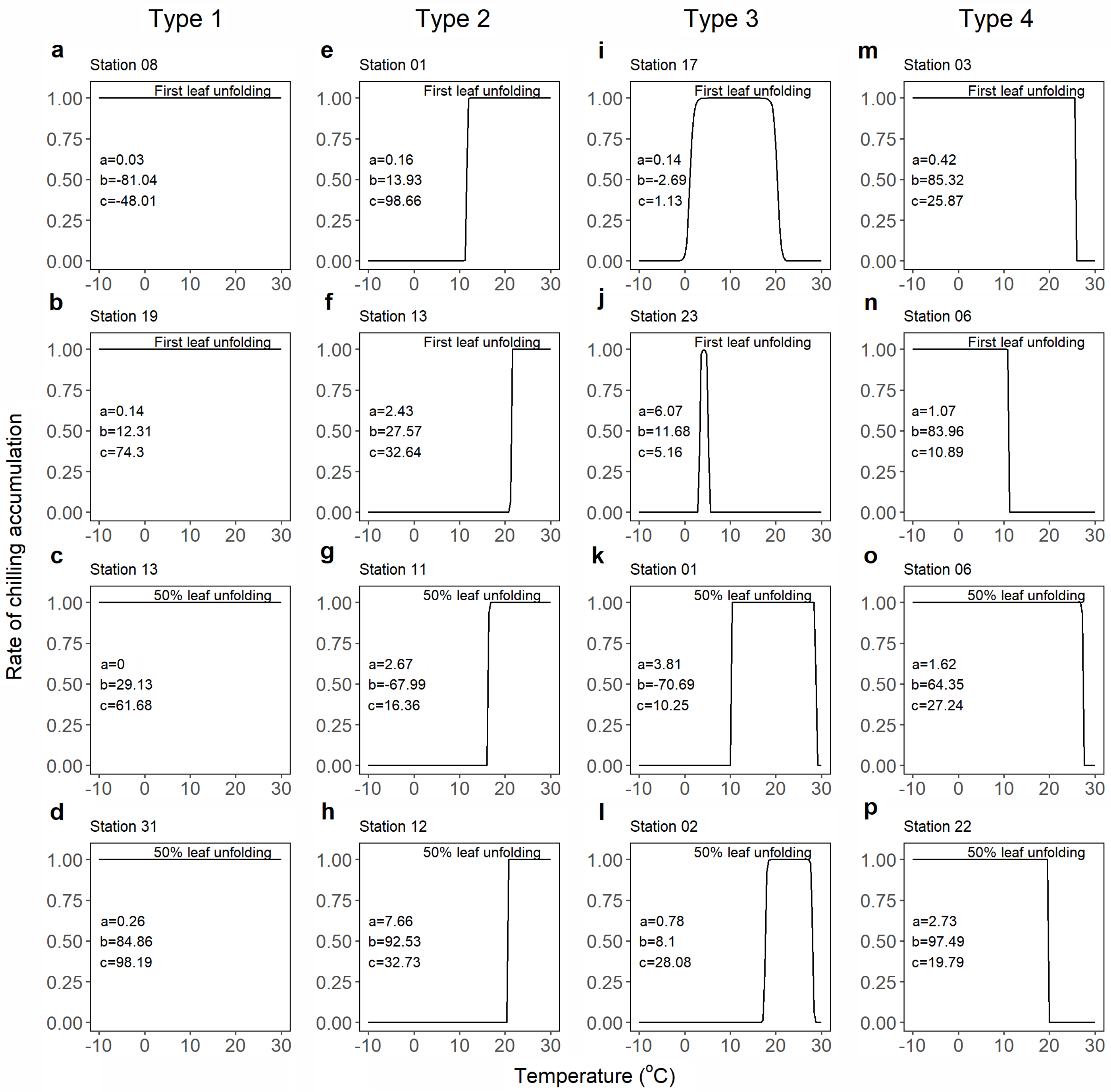

Type 1
Type 2
Type 3
Type 4
a
Station 08
First leaf unfolding
a=0.03
b=-81.04
c=-48.01
b
Station 19
First leaf unfolding
a=0.14
b=12.31
c=74.3
c
Station 13
50% leaf unfolding
a=0
b=29.13
c=61.68
d
Station 31
50% leaf unfolding
a=0.26
b=84.86
c=98.19
e
Station 01
First leaf unfolding
a=0.16
b=13.93
c=98.66
f
Station 13
First leaf unfolding
a=2.43
b=27.57
c=32.64
g
Station 11
50% leaf unfolding
a=2.67
b=-67.99
c=16.36
h
Station 12
50% leaf unfolding
a=7.66
b=92.53
c=32.73
i
Station 17
First leaf unfolding
a=0.14
b=-2.69
c=1.13
j
Station 23
First leaf unfolding
a=6.07
b=11.68
c=5.16
k
Station 01
50% leaf unfolding
a=3.81
b=-70.69
c=10.25
l
Station 02
50% leaf unfolding
a=0.78
b=8.1
c=28.08
m
Station 03
First leaf unfolding
a=0.42
b=85.32
c=25.87
n
Station 06
First leaf unfolding
a=1.07
b=83.96
c=10.89
o
Station 06
50% leaf unfolding
a=1.62
b=64.35
c=27.24
p
Station 22
50% leaf unfolding
a=2.73
b=97.49
c=19.79
Rate of chilling accumulation
Temperature (°C)
